# Discovery of CD80 and CD86 as recent activation markers on regulatory T cells by protein-RNA single-cell analysis

**DOI:** 10.1101/706275

**Authors:** Dominik Trzupek, Melanie Dunstan, Antony J. Cutler, Mercede Lee, Leila Godfrey, Dominik Aschenbrenner, Holm H. Uhlig, Linda S. Wicker, John A. Todd, Ricardo C. Ferreira

**Author notes:** Co-senior authors: Correspondence: Prof. John A. Todd:, Dr. Ricardo C. Ferreira.

## Abstract

The transcriptomic and proteomic characterisation of CD4^+^ T cells at the single-cell level has been performed traditionally by two largely exclusive types of technologies: single cell RNA-sequencing (scRNA-seq) technologies and antibody-based cytometry. Here we demonstrate that the simultaneous targeted quantification of mRNA and protein expression in single-cells provides a high-resolution map of human primary CD4^+^ T cells, and identified precise trajectories of Th1, Th17 and regulatory T-cell (Treg) differentiation in blood and tissue. Furthermore, the sensitivity provided by this massively-parallel multi-omics approach revealed novel insight into the mechanism of expression of CD80 and CD86 on the surface of activated CD4^+^ Tregs and demonstrate their potential to identify recently activated T cells in circulation. This transcriptomic and proteomic hybrid technology provides a cost-effective solution to dissect the heterogeneity of immune cell populations, including more precise and detailed descriptions of the differentiation and activation of circulating and tissue-resident cells in response to therapies and in stratification of patients.

## Background

Our understanding of the human immune system has been greatly influenced by the technological advances leading to the ability to precisely quantify mRNA and/or protein expression at the single-cell level. In particular, the implementation of flow cytometry as a routine and widely-accessible research tool has shaped much of our current knowledge about the complexity of the immune system. With increased availability of fluorochrome-conjugated antibodies and more powerful lasers, flow-cytometric assays allow typically 15-20 parameters that can be assessed in parallel. Developments in single-cell mass cytometry (CyTOF) have similarly allowed the simultaneous assessment of the expression of up to 50 protein targets using heavy metal-labeled antibodies [1].

The advent of single-cell RNA-sequencing (scRNA-seq) has provided an unprecedented opportunity to investigate the global transcriptional profile at the single-cell level. In contrast to cytometry-based technologies, which are limited to the concurrent detection of up to a few tens of protein markers, scRNA-seq technologies allow to profile the entire transcriptome. With the concomitant reduction in sequencing costs, there has been a recent explosion in scRNA-seq platforms available to immunologists [2, 3]. These fundamentally differ in the cell capture methods and resulting sensitivity, ranging from a few hundreds of cells profiled with high sensitivity, using plate-based capture methods such as SMART-seq2 [4], tens of thousands of cells with lower sensitivity using whole-transcriptome scRNA-seq platforms, such as 10X Genomics [5], Seq-Well [6] or Drop-seq [7].

Despite the growing popularity of whole-transcriptome scRNA-seq, two main issues still affect the performance of these platforms: cost and sensitivity. Even at high sequencing coverage, resulting in increased sequencing costs, stochastic dropout is a well-known issue of scRNA-seq, leading to an inflation of zero-expression measurements. Furthermore, although several methods have been developed to impute missing expression values, questions remain about the performance of these methods [8]. This technical limitation is particularly relevant for resting primary cells, such as CD4^+^ T cells, and mainly limits the robust detection and quantification of lowly expressed genes, including lineage-defining transcription factors, which are critical for cell-type identification and annotation. An important recent technical advance has been the development of new methods, such as CITE-seq [9] and REAP-seq [10], allowing the combination of whole-transcriptome scRNA-seq with measurement of protein expression at the single-cell level using oligo-conjugated antibodies. These methods provide critical insight into cell function and increased clustering resolution, although the resulting sequencing cost, especially when combining large numbers of antibodies targeting highly expressed proteins, still limits the use of this technology as a widely-applicable immunophenotyping tool.

In this study we describe an integrated targeted scRNA-seq workflow, to simultaneously quantify the expression of 397 genes at the mRNA level and up to 68 genes at the protein level with oligo-conjugated antibodies (AbSeq) [11]. We sought to assess the sensitivity and cost-effectiveness of this multi-omics system to immunophenotype human primary CD4^+^ T cells at the single-cell level, and to identify discrete cell-states providing potential new insight into the functional heterogeneity of T cells. By combining the expression of a targeted set of genes with the highly quantitative measurement of key protein markers, we have generated a high-resolution map of human CD4^+^ T cells in blood and tissue, and delineated distinct trajectories of T-cell differentiation associated with a gradient of activation, which were apparent even in resting primary cells. Our data also shows very clearly the frequent low correlation between mRNA and protein expression in primary CD4^+^ T cells, thereby challenging the current dogma that our current understanding of the immune system can be re-defined based on single-cell transcriptional data alone. These attributes provided novel evidence for the potential mechanisms leading to the expression of exogenous CD80 and CD86 on the surface of human primary Tregs, thus revealing a biomarker for activated Tregs that have recently contacted antigen presenting cells expressing these T-cell co-stimulatory signaling molecules.

## Results

### Simultaneous protein quantification increases the power of scRNA-seq to dissect the functional heterogeneity of human CD4^+^ T cells

In this study we wanted to investigate the power of a unified high-throughput experimental workflow combining targeted scRNA-seq and the quantification of protein expression at the single-cell level to dissect the heterogeneity of resting human primary CD4^+^ T cells in blood. To address this question, we initially profiled the expression of 397 genes at the mRNA level, coupled with 37 protein targets (**Table S1**) using the AbSeq technology [11] in CD4^+^ T cells isolated from blood of a systemic lupus erythematosus (SLE) patient. To enrich for the relative distribution of two less abundant CD4^+^ T-cell subsets: (i) CD127^low^CD25^+^ T cells, containing the Treg population; and (ii) CD127^low^CD25^low^ T cells, containing a subset of non-conventional CD25^low^FOXP3^+^ Tregs previously characterised in autoimmune patients [12], we devised a FACS-sorting strategy to isolate and profile equal numbers of cells from the three defined T-cell subsets (Fig. 1a). Following FACS sorting, cells from each subset were labeled with a barcoded oligo-conjugated antibody (sample tag) prior to cell loading – a method related to the recently described cell-hashing technique [13] – to identify their original sorting gate and to assess the frequency of cell multiplets obtained in this experiment.

**Fig. 1.**
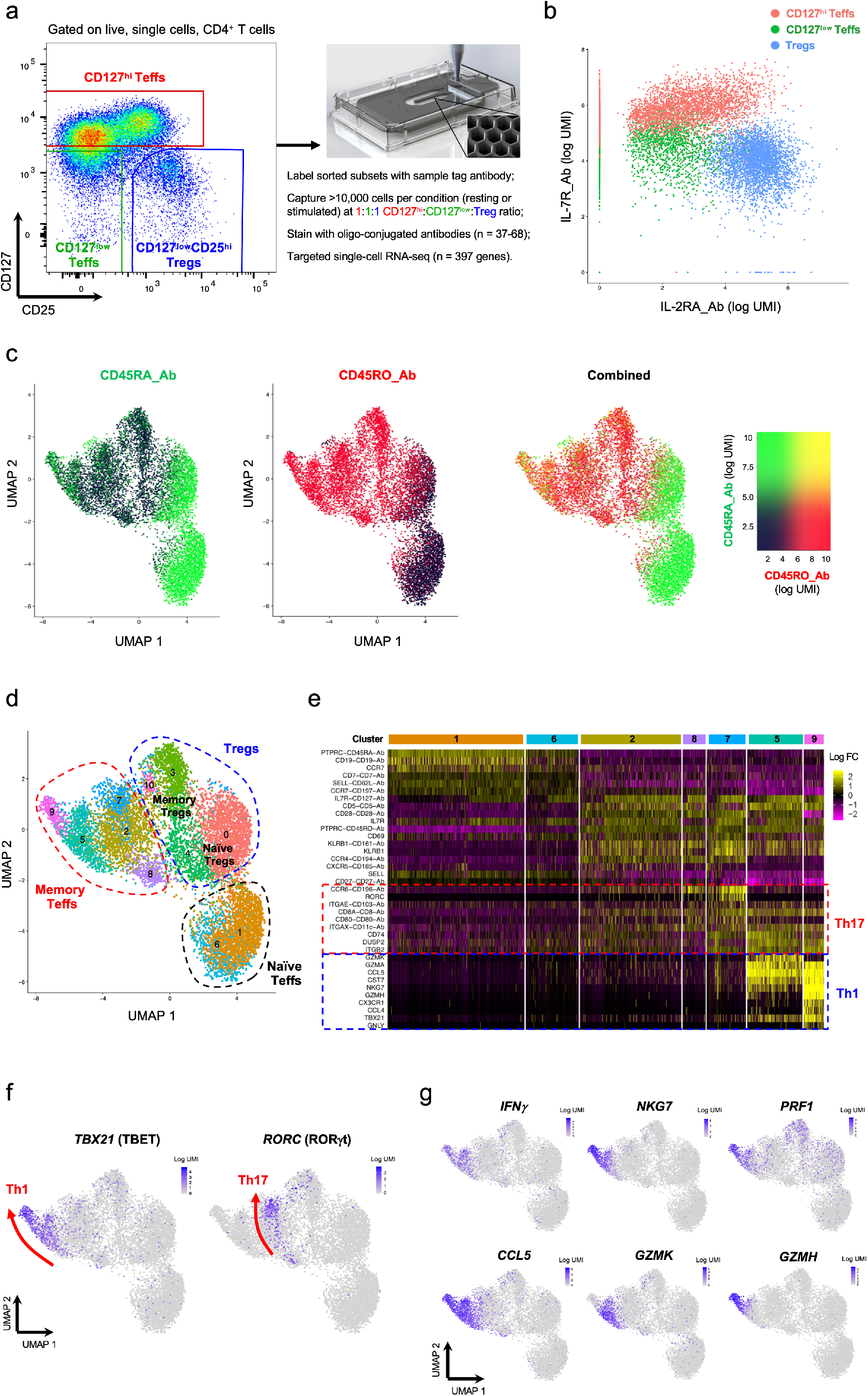
Combined single-cell transcriptional and proteomics immunophenotyping provides a high-resolution map of human primary CD4^+^ T cells in blood. (**a**) Summary of the experimental workflow. FACS plot depicting the sorting strategy for the isolation of the three assessed CD4^+^ T cell populations. (**b**) Two-dimensional plot depicting the expression of IL-7R and IL-2RA at the protein level using oligo-tagged antibodies (AbSeq) on each captured single cell. Cells are colored according to their respective sorting gate, as assessed using oligo-conjugated sample-tagging antibodies. (**c**) Uniform manifold approximation projection (UMAP) plot depicting clustering of all captured CD4^+^ single cells using the combined proteomics and transcriptomics data. Expression levels of the CD45RA (black to green) and CD45RO (black to red) isoforms using AbSeq antibodies are depicted in the plot. (**d**) UMAP plot depicting the clustering of resting primary CD4^+^ T cells (n = 9,708) isolated from blood of a systemic lupus erythematosus (SLE) patient. Identifying cluster numbers are assigned consecutively based on the total number of cells contained in each identified cluster. (**e**) Heatmap displaying the top 10 differentially expressed genes in each identified resting CD4^+^ Teff cluster. (**f**) UMAP plots depicting the expression of the CD4^+^ T-cell lineage-defining transcription factors TBET (Th1) and RORgt (Th17) in resting CD4^+^ T cells. Arrows recapitulate the identified axis of Th1 and Th17 differentiation, and are supported both on the gradient of expression of the respective lineage-restricted transcription factor (TBET and RORgt, respectively), but also on the developmental trajectories identified by pseudotime analysis depicted in Fig. 3. (**g**) Expression of the effector-type cytokine transcripts *IFNG*, *NKG7*, *PRF1*, *CCL5*, *GZMH* and *GZMK* in resting CD4^+^ T cells.

A total of 9,898 captured cells passed initial QC, of which a small proportion (1.9%; **Table S2**) were assigned as multiplets and excluded from the analysis. Of note, we observed complete sequencing saturation of the mRNA library, assessed as the number of cDNA molecules with a novel Unique Molecular Identifier (UMI) identified with increasing sequencing coverage, for a coverage of <3,000 reads/cell (Fig. S1a). In contrast, we obtained an 80% sequencing saturation at a coverage of >6,000 reads/cell for the AbSeq library (Fig. S1b). This is illustrated in the large dynamic range of expression of the protein targets, reaching up to thousands of unique copies in cells displaying higher levels of expression (Fig. S1c). Of note, the distribution of most proteins, including all those that are known not to be expressed on CD4^+^ T cells was centered around zero copies (Fig. S1c), which demonstrates the high specificity of the AbSeq immunostainings. To further test the sensitivity of the AbSeq protein measurements, we next generated a two-dimensional plot depicting the AbSeq expression of IL-2RA (CD25) and IL-7R (CD127), which we found to recapitulate the flow cytometric profile obtained with the same two markers used for the FACS-sorting of the assessed T-cell subsets (Fig. 1b). Furthermore, by overlaying the sample-tag information, we were able to confirm that the expression profiles of CD127 and CD25 mimicked the sorting strategy precisely for all three subsets (Fig. 1b), therefore illustrating the highly quantitative nature of the protein measurements.

Next we performed unsupervised hierarchical clustering combining the mRNA and protein expression data and visualised the clusters in a two-dimensional space using Uniform Manifold Approximation and Projection (UMAP) [14]. One of the main drivers of functional differentiation in CD4^+^ T cells is the acquisition of a memory phenotype in response to antigen stimulation, typically marked by the expression of CD45RA on naïve cells and CD45RO on memory cells. However, because these are splice forms of the same gene (*PTPRC*; CD45), its discrimination cannot be achieved using UMI-based scRNA-seq systems targeting the 3’ or 5’ ends of the transcript. By measuring the expression of the two isoforms at the protein level, we were able to identify a marked expression gradient associated with a gradual loss of CD45RA and concomitant gain of CD45RO along the first component of the UMAP plot, indicating that the acquisition of a memory phenotype is indeed the main source of biological variation driving the clustering of CD4^+^ T cells (Fig. 1c). One notable exception was the re-expression of CD45RA in the most differentiated memory cells (Fig. 1c). This observation is consistent with the phenotype of differentiated effector memory CD4^+^ T cells that re-express CD45RA (TEMRAs) [15], and illustrates the power of this highly multiparametric approach to identify subtle alterations in CD4^+^ T-cell states, while mitigating the potential issue of cell-type misclassification based on a few prototypical markers such as CD45RA/RO.

### Single-cell mRNA and protein immunophenotyping identifies distinct trajectories of CD4^+^ T cell differentiation in blood

Integration of the multiparametric transcriptional and proteomics data generated provided distinct clustering of CD4^+^ T cells into discrete clusters along the naïve/memory differentiation axis (Fig. 1d). We observed an increased number of clusters within the memory compartment, marked by the differential expression of defined sets of signature genes (Fig. 1e and Fig. S2a), which was consistent with the increased phenotypic diversity in differentiated CD4^+^ T-cell subsets. Moreover, we observed that the expression of the canonical Th1 (*TBX21*, encoding TBET) and Th17 (*RORC*, encoding RORgt) lineage-defining transcription factors was restricted to specific clusters within the effector memory T-cell (mTeff) population (Fig. 1f), indicating that these clusters are highly enriched for Th1 and Th17 Teffs. More importantly, we observed a distinct gradient of expression of these transcription factors. Consistent with this gradient of functional differentiation, we observed a marked co-expression of canonical Th1 effector-type molecules with the expression of TBET (Fig. 1f), revealing a subset of highly activated Th1 T cells with a putative cytotoxic profile in the blood of this SLE patient. Similarly, a gradient of expression of Th17-signature genes, including *RORC*, could be observed from cluster 8 to 7 (Fig. 1e,f), indicating a trajectory of Th17 differentiation.

In addition to resting CD4^+^ T cells, we also profiled the same subsets of cells following short *in vitro* stimulation (90 min) with PMA + ionomycin, to assess cell-type specific cytokine production. Similarly to the resting condition, *in vitro* stimulated CD4^+^ T cells formed discrete clusters along the naïve-memory differentiation axis (Fig. S3a,b). Furthermore, we observed a consistent induction of expression of Th1 (IFNg) and Th17 (IL-22) type cytokines that were restricted to the respective Th1 and Th17 clusters (Fig S3c,d). Furthermore, although primers for the Th2 transcription factor GATA-3 were not included in this assay, therefore precluding the annotation of Th2 cells in resting CD4^+^ T cells, we noted that *in vitro* stimulation revealed a distinct cluster of Th2 mTeffs cells marked by the expression of Th2-type cytokines, such as IL-13 (Fig S3e), but also IL-4, IL-5 and IL-9 (Fig. S2b).

Recently, several scRNA-seq studies have refined our understanding of the heterogeneity of CD4^+^ Tregs and their functional adaptation in tissues, in both mice and humans [16, 17]. Given the sorting strategy used in this study, we were able to significantly enrich our CD4^+^ T-cell dataset for the Treg population, which is highly enriched within the CD127^low^CD25^hi^ population. Consistent with this enrichment strategy, we identified a large Treg population marked by the expression of the transcription factor FOXP3, but also other classical Treg signature genes, including HELIOS (encoded by *IKZF2*), IL-2RA, CTLA-4 or TIGIT (Fig. 2a,b). In agreement with their Treg-specific transcriptional programme, we found a marked suppression of IL-2 transcription in Tregs following *in vitro* stimulation (Fig. S3f). Notably, we found that the differentiation of Tregs from a naïve to memory phenotype was strongly associated with the expression of two transcription factors: BACH2 and BLIMP1 (encoded by *PRDM1*). These two key transcription factors displayed a very distinct mutually exclusive expression pattern, with high expression of *BACH2* in naïve cells, declining gradually – with concomitant gradual increase in *PRDM1* expression – along the naïve-memory differentiation axis (Fig. 2c). The gradual increase of *PRDM1* expression was found to be strongly associated with the expression of Treg activation markers such as *HLA-DRA, DUSP4* and *CD39* (Fig. 2d), and revealed a trajectory of Treg activation in resting primary CD4^+^ T cells. These data suggest that the transcriptional interplay between BACH2 and BLIMP-1 is critical to regulate the differentiation of memory Treg subsets, which is in agreement with previous data in both mouse and humans [16]. The dynamic interplay of *BACH2* and *PRDM1* in the differentiation of Tregs was even more pronounced following *in vitro* stimulation (Fig. S4), which further supports the hypothesis that they are primary regulators of the transcriptional programme associated with the differentiation of suppressive activated Tregs in humans in response to antigen stimulation.

**Fig. 2.**
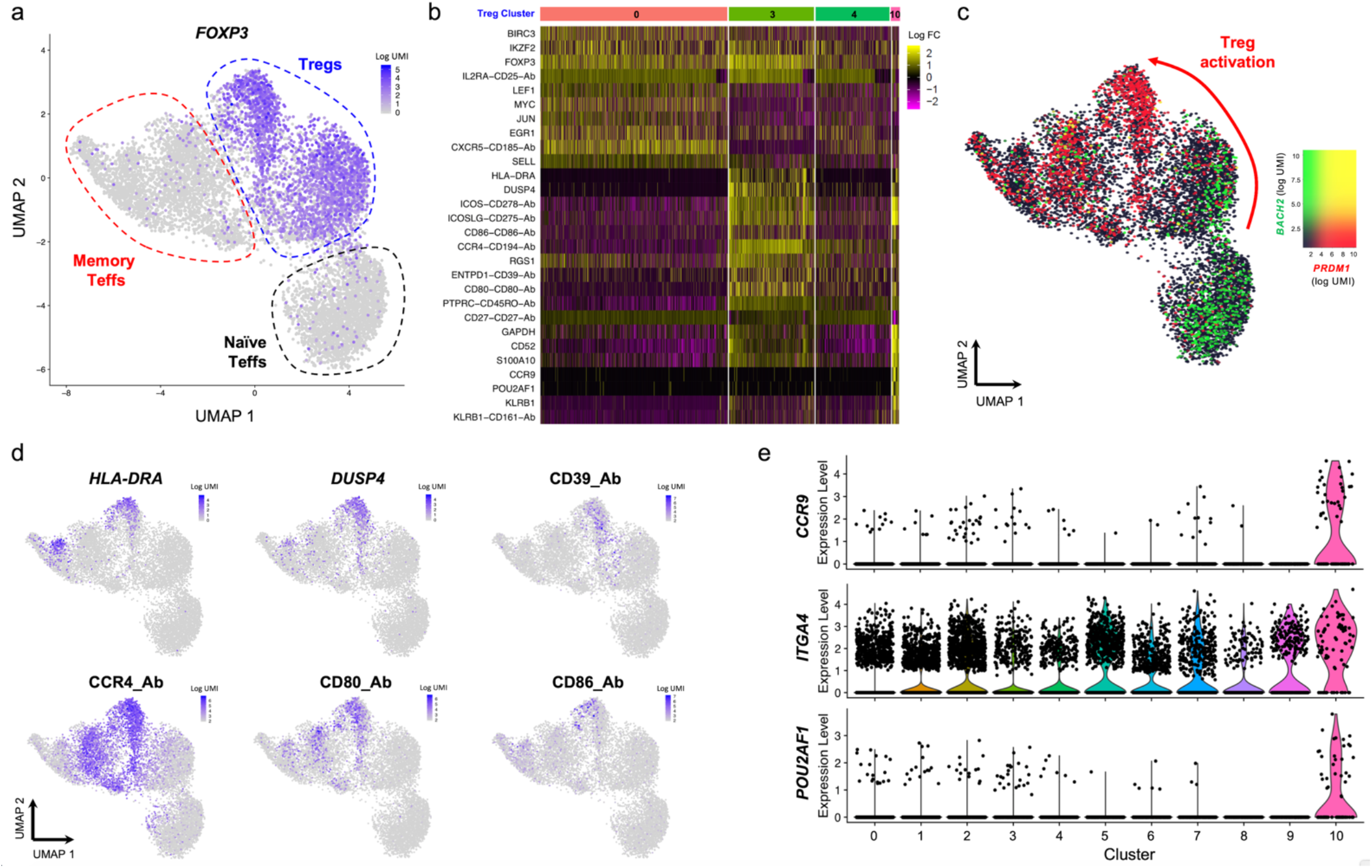
Integrated single-cell targeted multi-omics approach provides identifies a trajectory of human CD4^+^ regulatory T cell (Treg) activation. (**a**) UMAP plot depicting the expression of the canonical Treg transcription factor FOXP3 in the identified resting CD4^+^ T-cell clusters. (**b**) Heatmap displaying the top 10 differentially expressed genes within the four identified resting Treg clusters, depicted in Fig. 1D. (**c**) UMAP plot depicting the overlaid expression of two key CD4^+^ T-cell transcription factor *BACH2* (black to green) and *PRDM1* (encoding BLIMP-1; black to red). (**d**) Illustrative example of the expression of highly differentially expressed genes within the cluster of activated Tregs (cluster 3), including *HLA-DRA* and *DUSP4* at the mRNA level and CD39, CCR4, CD80 and CD86 at the protein level. (**e**). Violin plots depicting the expression of *CCR9*, *ITGA4* and the transcription factor *POU2AF1* in each defined CD4^+^ T-cell cluster.

Statistical methods are currently being developed to identify and reconstruct developmental trajectories from heterogeneous scRNA-seq datasets using pseudotime analysis. To validate our findings, we next applied the recently developed partition-based graph abstraction (PAGA) method [18] to reconstruct the developmental trajectories in our dataset. Consistent with our previous findings, pseudotime analysis revealed a gradient of T-cell differentiation along the naïve-memory differentiation axis, which lead to the identification of three distinct differentiation pathways associated with the acquisition of a Th1, Th17 or Treg phenotype (Fig. 3a,b). These identified differentiation trajectories were associated with gradual increased expression of the lineage-specific transcription factors TBET, RORgt and FOXP3, which regulate the transcriptional programme associated with the respective T-cell lineages (Fig. 3c-e). In particular, we confirmed a very distinct and gradual differentiation of the Th1 lineage in this SLE patient, leading to the temporal acquisition of activated Th1 cells expressing IFN-g in cluster 5 and the terminal differentiation of a subset with cytotoxic profile (cluster 9). Of note, this analysis identified cluster 2 as an intermediate memory Teff cell state, leading to the differentiation of either Th1 (cluster 5 and 9) or Th17 (cluster 7) T cells. Moreover, pseudotime analysis also recapitulated the acquisition of an activated Treg trajectory from naïve Tregs (cluster 0) to activated memory Tregs (cluster 3), which was regulated by the mutually exclusive expression of the BACH2 and BLIMP-1 transcription factors (Fig. 3e). An intriguing observation was the identification of cluster 8 representing a potential intermediate T-cell state on a Treg-Th17 developmental pathway, which is consistent with the plasticity and putative common co-evolutionary origin between these two lineages [19].

**Fig. 3.**
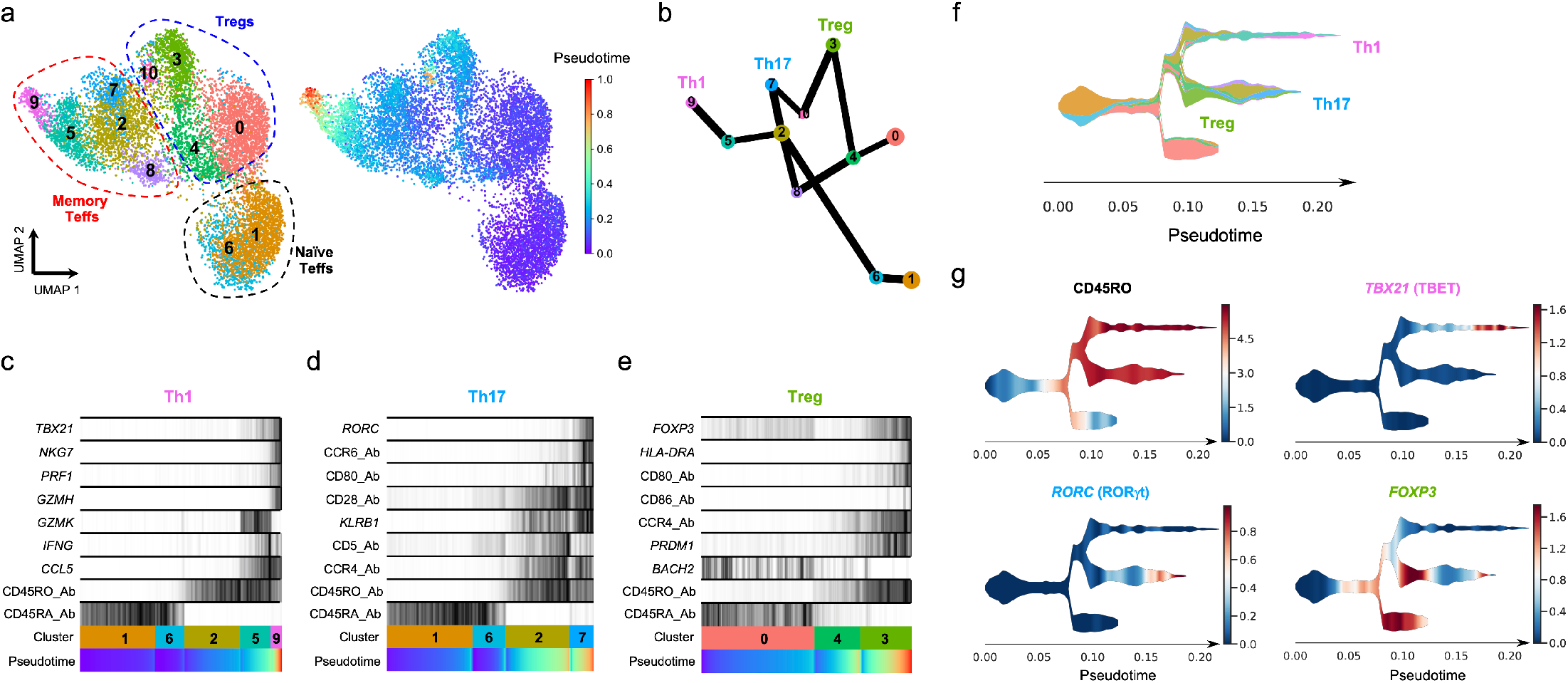
Pseudotime analysis reveals distinct trajectories of CD4^+^ T cell differentiation *in vivo*. (**a**) UMAP plots depicting the inferred diffusion pseudotime of each single-cell in the identified T-cell clusters. (**b**) Graph reconstructing the developmental trajectories between the identified T-cell clusters. Edge weights represent confidence in the presence of connections between clusters. The analysis was performed in the combined transcriptional and proteomics data using the Partition-based graph abstraction (PAGA) method. (**c-e**) Reconstructed PAGA paths for the differentiation of the identified Th1 (**c**), Th17 (**d**) and Treg (**e**) lineages. Expression of the lineage-specific transcription factors and selected differentially expressed genes is depicted for each trajectory. (**f**) Schematic representation of the identified lineage differentiation trajectories using the Single-Cell Trajectories Reconstruction (STREAM) method. Colour code corresponds to the cluster assignment depicted in panel A. (**g**) Expression of the memory-associated CD45RO isoform and the lineage-specific transcription factors *TBX21* (encoding TBET), *RORC* (encoding RORgt) and *FOXP3* is depicted along the identified developmental branches.

The identification of the temporal differentiation of these T-cell lineages was also recapitulated using Single-cell Trajectories Reconstruction (STREAM) [20], another method that has been developed to visualise developmental trajectories using multi-omics data (Fig. 3f). Further supporting a putative common developmental pathway of Treg and Th17 cells, STREAM analysis also identified FOXP3^+^ mTregs as a less differentiated T-cell state, which share a developmental trajectory with differentiated RORgt^+^ Th17 cells, with cluster 8 representing an intermediate transitional cell-state in this trajectory (Fig. 3g). These data illustrate not only the sensitivity of the targeted scRNA-seq approach to sensitively quantify lowly expressed transcription factor genes, but also highlight the power of this integrated multi-omics approach to identify subtle cell-state transitions. The large number of captured single-cells and the combination of protein and mRNA measurement that we obtained in this study makes this dataset particularly well suited to identify continuous cell-state transitions and reconstruct the differentiation trajectories from resting human primary T cells.

### Protein expression of CD80 and CD86 marks a subset of recently activated D4^+^ Tregs in circulation

A feature of the most activated Treg cluster (cluster 3), was the marked increased expression of CD80 and CD86 at the protein level (Fig. 2b,d), two T-cell costimulatory molecules usually expressed in antigen presenting cells (APCs). These findings were recapitulated on the pseudotime analysis, which identified CD80/CD86 protein expression as markers of the temporal Treg differentiation trajectory (Fig. 3e). Consistent with their APC-restricted expression pattern, we observed virtually no detectable expression of either *CD80* or *CD86* at the mRNA level in either resting or *in vitro* stimulated CD4^+^ T cells (Fig. 4a,b), suggesting a likely exogenous source of the detected protein expression.

**Fig. 4.**
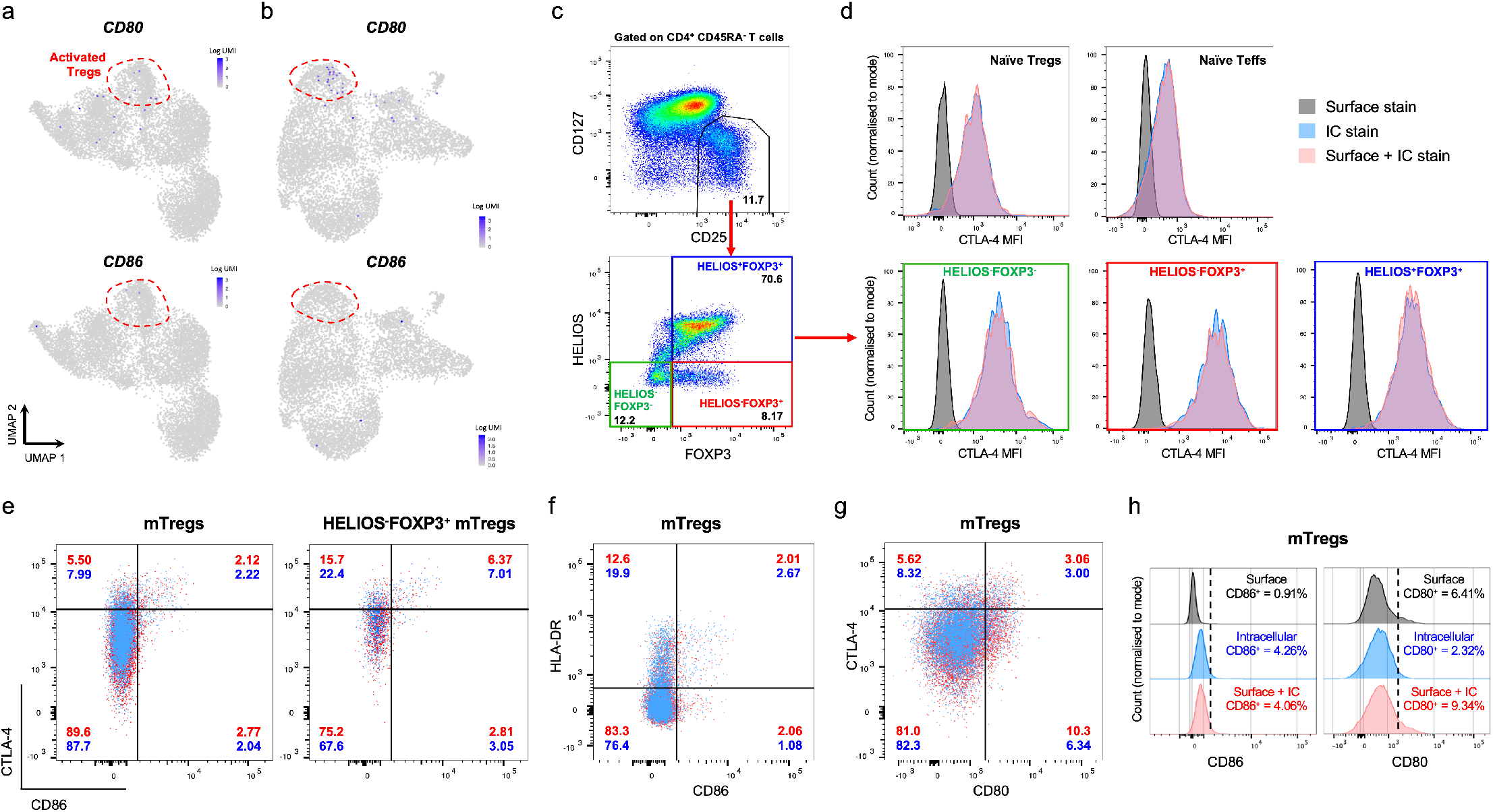
Surface expression of the co-stimulatory molecules CD80 and CD86 marks a subset of highly activated CTLA-4^+^ human Tregs *in vivo*. (**a,b**) UMAP plots depicting the mRNA expression levels of the co-stimulatory molecules CD80 and CD86 in the identified CD4^+^ T-cell subsets in resting (**a**) and *in vitro* stimulated (**b**) conditions. Dashed lines delineate the identified activated Treg clusters. (**c**) Gating strategy for the delineation of the Treg subsets according to the intracellular expression of the canonical Treg markers FOXP3 and HELIOS. (**d**) Histograms depicting the mean fluorescence intensity (MFI) of the CTLA-4 expression in the assessed CD4^+^ T cell subsets. CTLA-4, CD80 and CD86 immunostaining was performed under three different experimental conditions: (i) Surface immunostaining only (black); (ii) Immunostaining after cell permeabilization and fixation (IC; blue); and (iii) Both surface immunostaining and after cell permeabilization and fixation (red). (**e**) Dot plots depicting the co-expression of CTLA-4 and CD86 in CD45RA^-^ CD127^low^CD25^+^ Tregs (mTregs) and in the HELIOS^-^FOXP3^+^ mTregs. Flow cytometric data was generated from the intracellular immunostaining of CTLA-4 and CD86. (**f,g**) Two dimensional plots depicting the co-expression of CTLA-4 and CD80 (**f**) and HLA-DR and CD86 (**g**) in mTregs. Flow cytometric data was generated from the intracellular immunostaining of the assessed markers. (**h**) Histograms depict the mean fluorescence intensity (MFI) of CD80 and CD86 expression in mTregs. Data was obtained from the flow cytometric assessment of CD80 and CD86 by (i) Surface immunostaining only (black); (ii) Immunostaining after cell permeabilization and fixation (blue); and (iii) Both surface immunostaining and after cell permeabilization and fixation (red). Data shown in this figure was generated using freshly isolated PBMCs from two healthy donors recruited from the Oxford Biobank (depicted in red and blue, respectively). Cell frequencies from the respective donor are also indicated for each assessed population.

Recently, two mechanisms have been postulated to lead to the expression of APC-restricted proteins on the surface of activated T-cells, and Tregs in particular: (i) a CTLA-4 dependent mechanism (trans-endocytosis), whereby CTLA-4 expression on the surface of activated T cells can lead to the removal and subsequent endolytic degradation of CD80/CD86 molecules from the surface of APCs [21]; and (ii) a TCR-dependent mechanism (trogocytocis), whereby TCR engagement with the peptide-MHC complex on APCs can lead to the re-expression of APC-restricted molecules on the surface of activated T cells [22]. To validate the detected expression of CD80 and CD86 in human resting primary CD4^+^ T cells, and to shed light into the mechanisms of expression, we next designed a flow cytometry panel to investigate the expression of these proteins in CD4^+^ T cells isolated from blood of two healthy donors. We differentiated between surface and intracellular expression of the assessed markers by performing immunostaining either prior or following cell fixation and permeabilisation. Consistent with this strategy, we confirmed that although the expression of CTLA-4 on the surface of CD4^+^ T cells is very low, intracellular staining revealed much higher expression, most notably in the different Treg subsets (Fig. 4c,d). Notably, we detected a remarkable co-expression between CD86 and CTLA-4. In fact, CD86 expression was only detected on CTLA-4^hi^ T cells, and was therefore mainly restricted to memory Treg (mTreg) subsets, on the cells expressing the higher levels of CTLA-4 (Fig. 4e). In addition, we also observed a strong co-expression of CD86 and HLA-DR, another marker associated with Treg activation (Fig. 4f). In contrast to CD86, the co-expression between CD80 and CTLA-4 on mTregs was less pronounced, although still displaying preferential expression on CTLA-4^hi^ cells (Fig. 4g). Furthermore, we also observed notable differences in the cellular location of CD80 and CD86 expression. While CTLA-4 and CD86 displayed a predominantly intracellular location, expression of CD80 was mostly restricted to the surface of the cells and was detected at similar levels using either surface or intracellular immunostaining (Fig. 4h). Taken together, these data reveal novel insight into the specific mechanisms of CD80 and CD86 protein expression on the surface of human Tregs, and suggest that they could be useful as novel biomarkers of recently activated T cells in the circulation and indicative of their engagement with CD80/CD86-expressing APCs.

### Multi-omics immunophenotyping approach identifies a rare subset of circulating CCR9^+^ Tregs displaying hallmarks of tissue-resident T cells

Another example of a rare T-cell population that we were able to identify in circulation using this multimodal immunophenotyping strategy was a subset of FOXP3^+^ T cells marked by the specific expression of the small intestine-homing chemokine receptor CCR9 (cluster 10; Fig. 2e). This cluster was also marked by increased expression of other classical markers of tissue residency and migration to the gut, such as *ITGA4* (CD49d) and *ITGAE* (CD103; Fig. 2e). One distinguishing feature of this subset was the expression of the transcription factor *POU2AF1* (Fig. 2e). Although *POU2AF1* (encoding OCA-B), has been mainly characterised as a B-cell specific transcription factor in blood, where it plays a role in B-cell maturation [23], it has also been recently shown in mice to regulate the maintenance of memory phenotype and function in previously activated CD4^+^ T cells [24], and the differentiation of T follicular helper (Tfh) cells in the tissue [25]. In agreement with this putative tissue-resident phenotype, pseudotime analysis demonstrated that these cells correspond to a highly differentiated cell state (Fig. 3a). These data suggest that this subset of CCR9^+^ T cells may represent a previously uncharacterised population of recirculating tissue-resident Tregs in humans, and provides an illustrative example of the power of this multiparametric immunophenotyping approach to identify rare immune populations and reveal novel insight into the biology of CD4^+^ T cells.

### Single-cell comparison of mRNA and protein expression levels reveals modest and variable levels of correlation in primary CD4^+^ T cells

Given that the main advantage of this combined targeted scRNA-seq and proteomic approach is the ability to immunophenotype large numbers of cells from multiple donors, we next investigated whether we were able to integrate data generated from independent experiments. We replicated the initial experiment using the same pre-sorting strategy to isolate the three assessed CD4^+^ T-cell subsets from an individual with type 1 diabetes and one healthy donor. To further test the potential of the protein quantification, we extended the AbSeq panel to 43 protein targets expressed on CD4^+^ T cells (**Table S1**). In agreement with the initial experiment, unsupervised clustering of the 23,947 cells passing QC revealed a similar discrimination of CD4^+^ T-cell subsets (Fig. 5a). More importantly, we found very good alignment of the data from the three donors, with minimal evidence of significant experimental batch effects (Fig. 5b). Analysis of the donor-specific distribution of the identified CD4^+^ T-cell clusters also showed that the frequency of the putative CD4^+^ cytotoxic Th1 subset (cluster 11), marked by the co-expression of TBET and effector-type cytokines, was highly increased in the SLE patient (Fig. 5c-e). To avoid age-specific differences in the relative distribution of the CD45RA^+^ naïve and CD45RO^+^ memory compartments in these donors, we normalised the analysis to the memory T-cell clusters only, which we were able to robustly annotate using the AbSeq data for the expression of CD45RA and CD45RO. Although we detected a few cells with this activated Th1 phenotype in circulation from every donor, there was a very substantial expansion in the SLE patient (2.5% of memory CD4^+^ T cells compared to 0.3% and 0.1% in the T1D patient and healthy donor, respectively; Fig. 5d), suggesting that it could represent a pathogenic CD4^+^ T-cell subset associated with systemic autoimmunity in this patient.

**Fig. 5.**
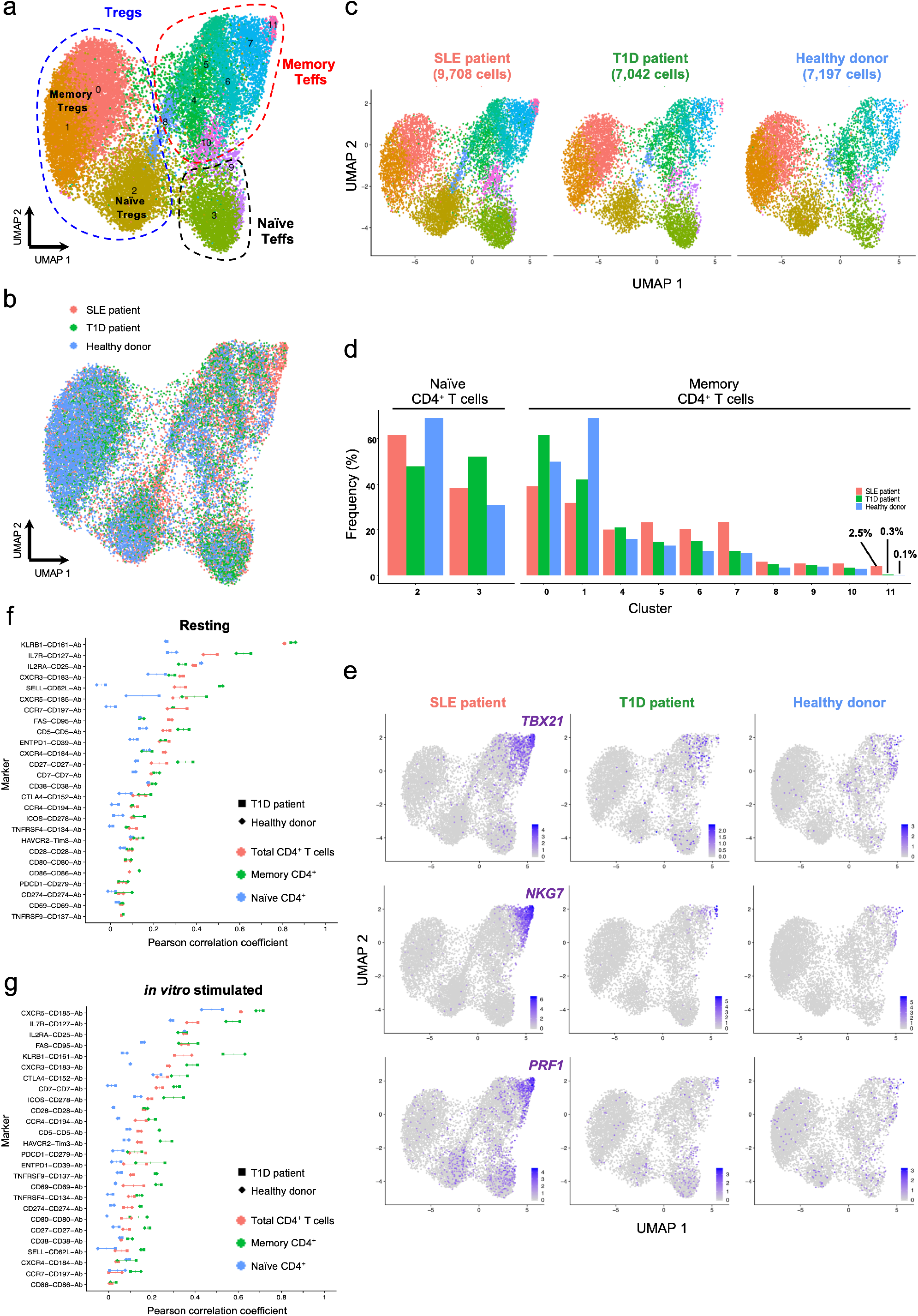
Data from independent experiments can be robustly integrated. (**a**) UMAP plot depicting the clustering of resting primary CD4^+^ T cells from one systemic lupus erythematosus (SLE; n = 9,708 cells) patient, one type 1 diabetes (T1D; n = 7,042 cells) patient and one healthy donor (n = 7,197 cells). Data was integrated from two independent experiments using the same CD4^+^ T-cell FACS sorting strategy (described in Fig. 1A). (**b**) Alignment of the integrated targeted transcriptomics and proteomics data generated from the three assessed donors in two independent experiments. (**c**) UMAP plots depicting the donor-specific clustering of the CD4^+^ T cells. (**d**) Relative proportion of the identified CD4^+^ T-cell clusters in each donor. Frequencies were normalised to either the annotated naïve or memory compartments to ensure higher functional uniformity of the assessed T-cell subsets and to avoid alterations associated with the declining frequency of naïve cells with age. (**e**) UMAP plots depicting the relative expression of the canonical Th1 transcription factor *TBX21* (encoding TBET) and the effector cytokines *NKG7* and *PRF1* on the three assessed donors. (**f, g**) Correlation (Pearson correlation coefficient) between mRNA and protein expression for 26 markers with concurrent mRNA and protein expression data in resting (**f**) and *in vitro* stimulated (**g**) CD4^+^ T cells. Correlation was calculated in total CD4^+^ T cells (red) or in the CD45RA^-^ memory (green) or CD45RA^+^ naïve (blue) T-cell subsets separately. Individual-level correlation in the type 1 diabetes (T1D) patient (square) and heathy donor (diamond) and median correlation in both donors are displayed in the figure.

The parallel quantification of mRNA and protein expression for a large number of genes expressed in CD4^+^ T cells in this experiment provided a unique opportunity to investigate their systematic correlation at the single-cell level. From the 43 proteins quantified with AbSeq, 26 were also assessed at the transcriptional level and detected in our CD4^+^ T-cell dataset. Generally, we observed relatively weak (mean Pearson correlation coefficient = 0.214) but variable levels of correlation in total resting CD4^+^ T cells, ranging from 0.049 for *TNFRSF9* to 0.808 in *KLRB1* (encoding CD161; Fig. 5f). Furthermore, we note that with the exception of *CXCR5*, the estimated correlations were very consistent between the two independent donors (Fig. 5f). These findings were consistent with previous observations [9, 10], and suggest that primary CD4^+^ T cells are highly specialised cells, where transcription is subject to tight regulation to avoid excessive energy consumption by the cell and to control effector function. As expected, by normalizing our analysis to a functionally more homogeneous population of memory CD4^+^ T cells, we observed higher levels of correlation (mean = 0.233), which is consistent with their increased expression of the majority of the assessed T-cell markers. A slightly decreased correlation (mean = 0.178) was observed in *in vitro* stimulated CD4^+^ T cells, suggesting an increased variance of protein and mRNA expression in activated CD4^+^ T cells (Fig. 5g).

### Parallel mRNA and protein profiling provides increased cell-type resolution of the heterogeneous CD45^+^ immune cell population in blood and tissue

To investigate how this combined targeted scRNA-seq and transcriptomics approach performs on a more heterogeneous population of immune cells, we isolated total CD45^+^ cells isolated from blood and a matching duodenal biopsy from two coeliac disease (CD) patients with active disease. In this experiment we captured 31,907 single-cells that passed QC and expanded the AbSeq panel to the detection of 68 protein targets (**Table S1**). As expected, we observed a very defined clustering of the different cell populations representing the CD45^+^ immune cells (Fig. 6a). Consistent with previous data [26, 27], we found a clear separation in cells isolated from either blood or the small intestine (Fig. 6b), indicating a strong transcriptional signature of tissue-residency. Furthermore, clustering of the cells isolated from either blood (Fig. 6c and Fig. S5a) or tissue (Fig. 6d) separately, revealed clear identification of the expected cell populations. The main distinction was the relative distribution of the main immune populations, with a marked increased representation of B-cell, NK-cell and CD14^+^CD16^-^ monocyte populations in blood, and a significant increase in the frequency of plasma cells in the small intestine. In agreement with our findings in CD4^+^ T cells, we found that the acquisition of a memory phenotype was the main driver of the clustering of both CD4^+^ and CD8^+^ T cells (Fig. S5b,c). In addition, we identified other clusters of non-conventional T cells, including a subset of gd T cells and mucosal-associated invariant T cells (MAIT) in blood, which shared similarities with the transcriptional signature of memory CD8^+^ T cells, marked by the expression of effector-type cytokines genes, such as *NKG7* (Fig. S5c). In contrast, tissue-resident CD4^+^ T cells isolated from the small intestine were restricted to a memory phenotype and displayed a markedly different subset distribution, including a substantially enlarged population of FOXP3^+^ Tregs (Fig. 6e,f). Moreover, the simultaneous assessment of the protein expression of CXCR5, ICOS and PD-1, identified a cluster of Tfh cells (Fig. 6e), which could be distinctly clustered along a trajectory of Tfh-cell activation, as illustrated by the gradient of expression of key Tfh functional transcripts, such as *IL21*, *CXCL13* and *BTLA* (Fig. 6g).

**Fig. 6.**
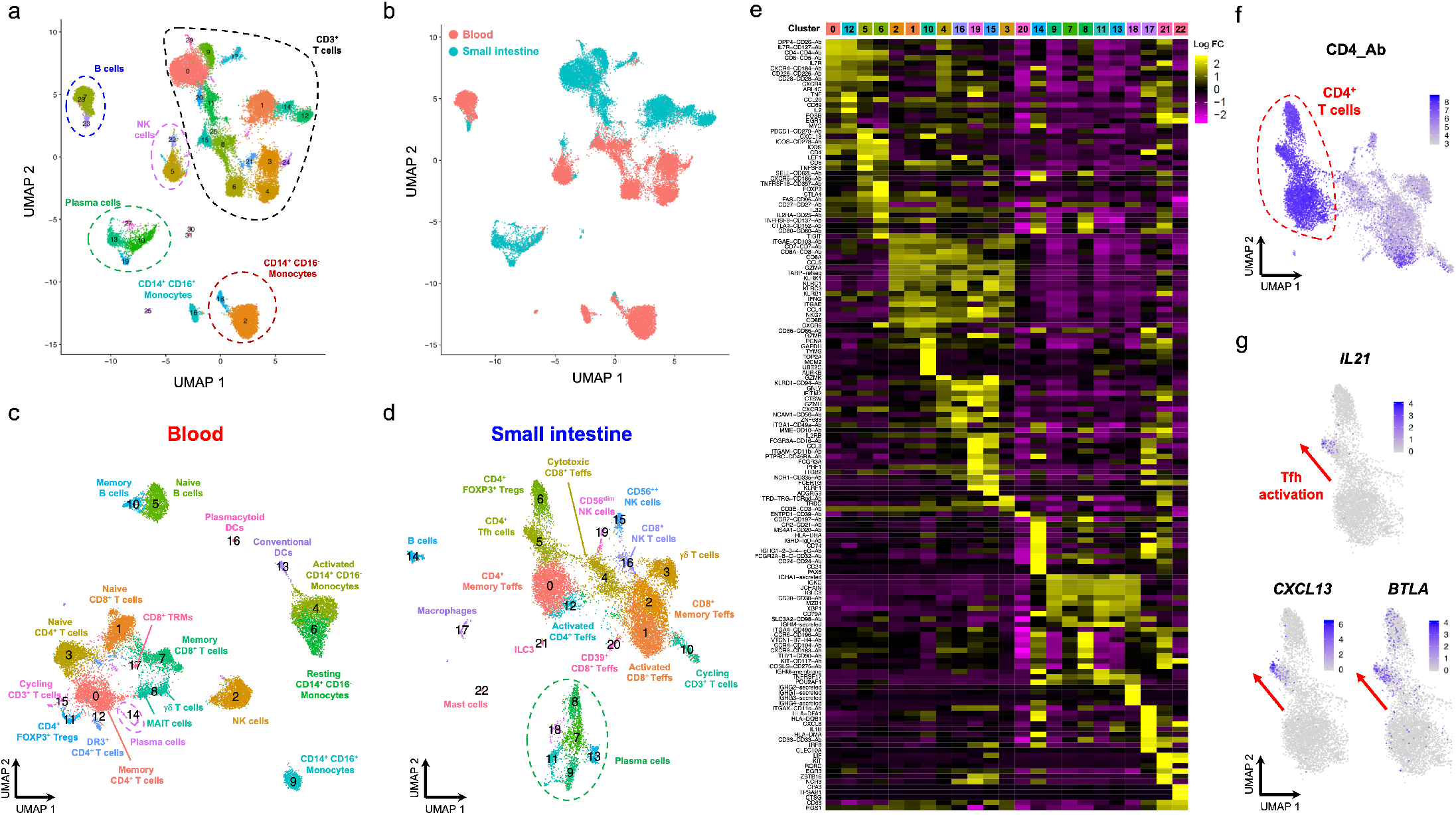
Targeted scRNA-seq and proteomics approach delineates distinct functional subsets in a heterogeneous CD45^+^ immune cell population isolated from blood and tissue. (**a**) UMAP plot depicting the clustering of the targeted scRNA-seq and transcriptional data of a heterogeneous population of total CD45^+^ cells (n = 31,907) isolated from blood and a paired duodenal biopsy from two coeliac disease (CD) patients with active disease. (**b**) Sample tag information identifies samples isolated from blood (red) or from the paired duodenal biopsy (teal). (**c, d**) UMAP plot depicting the clustering of the CD45^+^ cells isolated from blood (**c**) or the paired duodenal biopsy (**d**). (**e**) Heatmap displaying the top 10 differentially expressed genes in each identified cluster from the CD45^+^ immune cells isolated from the duodenal biopsies. (**f**) UMAP plot depicting the expression of the CD4 at the protein level (AbSeq) within the CD3^+^ T cells isolated from the small intestine. (**g**) Gradient of expression of the Tfh effector genes *IL21*, *CXCL13* and *BTLA* in tissue-resident CD4^+^ T cells. DR3, death-receptor 3 (encoded by *TNFRSF25*); TRM, tissue-resident memory T cells; MAIT, mucosal-associated invariant T cells; ILC3, type 3 innate lymphoid cell.

Similarly, we also identified distinct trajectories of cell differentiation in other immune cell types, as illustrated by the gradient of differentiation and class-switching of B cells in blood (Fig. S6a-c). Peripheral B cells were clearly dominated by a naïve IgD^+^IgM^+^ CD27^-^ subset, and only a small fraction of class-switched IgG^+^ CD27^+^ memory B cells, which was consistent with the young age of the CD patients. In contrast, tissue-resident B cells were much less abundant and contained mostly cells with a class switched IgG^+^ CD27^+^ memory phenotype. In addition, we identified a vastly expanded population of antibody-secreting plasma cells (Fig. S6d,e). Of note, because we were able to specifically assess the expression of the secreted Ig isotypes, we could also discriminate precisely the different functional plasma cell subsets, including a very abundant population of IgA-secreting plasma cells (Fig. S6f), which are known play a critical role in interaction with the microbiome in the gut. Together, these data provide an example of the power of this multimodal approach to identify trajectories of cell differentiation and cell states in diverse immune cell and tissue types.

## Discussion

The advent of scRNA-seq has proved to be a transformative technology that is shaping our understanding of the complexity and function of the human immune system [28, 29]. However, currently, both the elevated costs to perform these experiments, as well as the reliance on transcriptional data alone, pose significant challenges to the widespread practical applicability of these technologies. In this study, we present an integrated, cost-effective, approach to sensitively assess simultaneous expression of mRNA and protein for hundreds of key immune targets at the single-cell level using the AbSeq technology [11].

Recently, two similar approaches, CITE-seq [9] and REAP-seq [10], have been described to measure protein expression using oligo-conjugated antibodies in parallel with scRNA-seq data. Furthermore, other applications are currently being developed to integrate the growing portfolio of single-cell omics technologies [30, 31]. A fundamental difference with the approach described in this study is that these technologies all rely on whole-transcriptome data, providing a high-level cross-sectional representation of all polyA mRNA transcripts in the cell. In contrast, by using targeted scRNA-seq, we are relying on prior knowledge to specifically assess the expression of hundreds of selected genes in single cells. Moreover, a targeted approach provides a more sensitive quantification of expression of the selected genes at a fraction of the cost to generate the sequencing libraries, as it avoids the detection of highly expressed invariant housekeeping genes, which take up the vast majority of the whole-transcriptome scRNA-seq libraries. The increased sensitivity of a targeted approach is particularly relevant for the accurate assessment of lowly-expressed genes with critical regulatory function, such as transcription factors, which can be poorly quantified using traditional whole-transcriptome scRNA-seq data. It therefore, provides a knowledge-based approach to validate and extend whole-transcriptome scRNA-seq findings, that can be widely implemented in any research or clinical setting. Similarly to other widely-implemented knowledge-based single-cell immunophenotyping tools such as flow-cytometry and CyTOF, the highly customisable nature of this approach is critical to investigate specific research questions with very high sensitivity and in larger number of samples. However, in contrast to CyTOF which is inherently time-consuming and requires the availability of large numbers of cells to maximise the information generated by each run, this technology is ideally suited for unique and highly valuable clinical samples, for which cell availability and number are major practical constraints. Furthermore, the digital nature and lack of spectral overlap issues with the AbSeq measurements provide a superior specificity and sensitivity compared to flow cytometry for the quantification of lowly-expressed proteins, allowing accurate detection of zero or very low copy numbers, which are usually difficult to discriminate by flow cytometry. A good example was the sensitive quantification of the co-stimulatory protein CD80 by AbSeq, whose expression was found to be restricted to activated T cells. In comparison, flow cytometric assessment of CD80 expression was much less well-resolved with higher background, resulting in lower dynamic range of expression comparing to AbSeq.

<u>The increased sensitivity and high number of parameters simultaneously assessed using this multi-omics approach can also lead to unanticipated novel biological findings. An illustrative example of this potential is the identification of the APC-restricted CD80 and CD86 co-stimulatory proteins as markers of activated Tregs in peripheral blood. Although mechanisms of CD80/86 protein expression on the surface of T cells, including trogocytosis and trans-endocytosis, have been previously demonstrated, they have been restricted to</u> either mouse models or to *in vitro* cell line models [32–34]. Our data demonstrate that expression of CD80 and CD86 at the protein level can also be detected in humans on the surface of primary T cells isolated *ex vivo* from blood, and reveal specific differences between the expression of these two proteins that could provide novel insight into the functional mechanism(s) leading to their expression. Notably, while CD86 expression was remarkably restricted to CTLA-4^hi^ Tregs and intracellular, CD80 was less dependent on CTLA-4 expression and mostly detected on the surface of T cells. Consistent with these results, the AbSeq data revealed that CD86 expression was restricted to a small subset of memory Tregs displaying hallmarks of a highly activated phenotype, supporting the hypothesis that CD86 molecules captured by CTLA-4 could be transiently expressed on the surface of activated Tregs prior to internalisation and degradation. In contrast to CD86, CD80 protein expression could also be detected in Tregs with lower CTLA-4 levels, and displayed broader expression profile in other activated T-cell subsets. In particular the pseudotime analysis in our dataset, identified CD80 as a marker of the temporal differentiation of Th17 cells, which may provide a mechanistic rationale for the recently reported suppression of Th17 differentiation in response to anti-CD80 treatment in mice [35]. Furthermore, we also note a distinct co-expression of CD80 protein and HLA-class II mRNA (*HLA-DRA*) in a subset of activated Th1 cells, which could indicate recent activation in the context of strong TCR signaling required to induce the differentiation of Th1 cells [36, 37].

Owing to the strong upregulation of CTLA-4 on activated T cells, our data cannot provide conclusive evidence for the occurrence of trans-endocytosis in primary human T cells. Nevertheless, the co-expression between CD86 and CTLA-4 and its restricted expression on activated Tregs, suggest that likely both trogocytosis and trans-endocytosis contribute to the expression of APC molecules on the surface of human primary CD4^+^ T cells. In addition, we also cannot rule out that the possibility that CD80 and CD86 are specifically expressed on activated CD4^+^ T cells, but that the mRNA has a very short half-life, a hypothesis that has been previously reported [38]. Nevertheless, these data point to subtle, previously uncharacterized, differences in the expression of CD80 and CD86 in different T cell subsets. To our knowledge, these findings provide the first evidence for the occurrence of these mechanisms in human primary T cells, and indicate that surface expression of CD80 and CD86 could be valuable new markers for the isolation of antigen-specific T cells for functional assays.

Another example of the of the potential of this multi-omics approach to reveal novel biological findings was the identification of a Th1 CD4^+^ T-cell subset with marked cytotoxic profile that we found to be selectively expanded in an SLE patient. Of interest, this subset displayed a phenotype that is consistent with a population of CXCR3^+^ PD-1^+^ CD4^+^ T cells that were recently found to be expanded in blood from SLE patients [39]. Although our current dataset is limited in patient numbers, the transcriptional profile of the heterogeneous CXCR3^+^PD-1^+^ CD4^+^ T-cells described by Caielli *et al* display the same distinct Th1 profile, marked by the expression of the transcription factor TBET and numerous cytotoxic effector molecules. Our data suggest that a specific enrichment with the cytotoxic CD4^+^ T-cell subset described in this study could be responsible for the reported enrichment of CXCR3^+^PD-1^+^ CD4^+^ T-cells in SLE patients, and provides additional functional characterisation of this heterogeneous subset at the single-cell level, which could be critical to further investigate its role in the pathogenesis of SLE.

An important finding from this study and other related studies [9, 10], is the generally low levels of correlation between mRNA and protein expression in primary CD4^+^ T cells at the single-cell level. One possible explanation for this observation is that reduced sensitivity of scRNA-seq to quantify mRNA expression, may be leading to an underestimation of the correlation coefficients. However, we note that there are notable exceptions, such as CD161, which displayed a high correlation between mRNA and protein levels at 0.847 in memory CD4^+^ T cells, demonstrating that a systematic error in the quantification of mRNA levels by scRNA-seq technologies is not the only factor contributing to the observed low level of correlation. These findings therefore underscore the importance of parallel protein quantification to better identify stable cellular phenotypes associated with cell function. In contrast to mRNA expression, proteins display a much larger dynamic range of expression and longer half-life [40, 41], resulting in much higher copy numbers, and a more accurate and reliable quantification compared to their mRNA counterparts. This is particularly relevant in differentiated resting primary cells, such as CD4^+^ T cells, where transcription is tightly regulated to maintain effector function. These low copy numbers result in increased stochastic variation in mRNA quantification and dropout rate, which impair the accuracy single-cell methods that rely only on transcriptional data. Furthermore, mRNA profiling provides only a snapshot into the current functional state of the cell, which can be better assessed with combined protein expression data. An illustration of the power of this combined multimodal approach is the detailed trajectories of differentiation that we identified in resting primary CD4^+^ T cells, which were recapitulated by precise gradients of mRNA expression. The sensitivity of these measurements combined with the high numbers of cells analysed lend themselves to identify gradual and subtle changes in cell states, which are critical to identify dynamic changes reflecting mechanisms of functional adaptation in a heterogeneous cell population.

In summary we here show that combined targeted scRNA-seq and protein expression analysis provides a high-resolution map of human immune cells in blood and tissue, and reveals novel biological insights into the biology of CD4^+^ T cells, as illustrated by the identification of CD80/CD86 expression on activated Tregs in circulation and the functional characterisation of a potential pathogenic cytotoxic CD4^+^ T-cell subset in SLE. Our data provide a proof-of-principle for the implementation of this integrated approach as a widely applicable, and cost-efficient research tool for immunologists that could be particularly valuable in a clinical setting for the characterisation of rare patient samples with limited cell numbers, as well as to assess the functional consequence at the single-cell level of targeting key biological pathways *in vivo*, in patients treated with immunotherapeutic drugs.

## Methods

### Subjects

Study participants included one SLE patient (37 y/o female), recruited from the Cambridge BioResource, and one T1D patient (16 y/o male) and one autoantibody negative healthy donor (14 y/o male) recruited from the JDRF Diabetes–Genes, Autoimmunity and Prevention (D-GAP) study.

Comparison of total CD45^+^ immune cells isolated from paired blood and a duodenal biopsy was performed in cells isolated from two paediatric coeliac disease (CD) patients with active disease (one 5 y/o male with Marsh scale disease score of 3c, and one 15 y/o male with Marsh scale disease score of 3b).

Flow cytometric assessment of the expression of CTLA-4, CD80 and CD86 in the Treg subsets was performed in two adult healthy donors (46 y/o female and 51 y/o male), recruited from the Oxford Biobank.

### Cell preparation and FACS sorting

T-cell centric assays were performed on cryopreserved peripheral blood mononuclear cells (PBMCs). Cryopreserved PBMCs were thawed at 37°C and resuspended drop-by-drop in X-VIVO (Lonza) with 1% heat-inactivated, filtered human AB serum (Sigma). Total CD4^+^ T cells were isolated by negative selection using magnetic beads (StemCell Technologies), and incubated with Fixable Viability Dye eFluor 780 (eBioscience) for 15 min at room temperature. After washing in PBS with 0.02% BSA cells were stained in 5ml FACS tubes (Falcon) with the fluorochrome-conjugated antibodies used for cell sorting and the BD AbSeq oligo-conjugated antibodies (BD Bioscience), according to the manufacturer’s instructions.

Cell sorting was performed using a BD FACSAria Fusion sorter (BD Biosciences) at 4°C into 1.5 mL DNA low bind Eppendorf tubes containing 500ul of X-Vivo with 1% heat-inactivated, filtered human AB serum. Following cell sorting, the three assessed T-cell subsets were incubated with sample tag antibodies (Sample multiplexing kit; BD Bioscience), washed 3 times in cold BD sample buffer (BD Biosciences) and counted. Samples were then pooled together in equal ratios in 620 ul of cold BD sample buffer at the desired cell concentrations – ranging from 20 to 40 cells/ul for an estimated capture rate of 10,000-20,000 single-cells – and immediately loaded on a BD Rhapsody cartridge (BD Biosciences) for single-cell capture.

For the *in vitro* stimulated condition, sorted CD4^+^ T-cell subsets were incubated in round-bottom 96-well plates (20,000 cells/well) at 37°C for 90 min in X-Vivo with 5% heat-inactivated, filtered human AB serum with a PMA and ionomycin cell stimulation cocktail (eBioscience), in the absence of protein transport inhibitors. Cells were harvested into FACS tubes, washed with cold BD sample buffer and further incubated with the BD AbSeq oligo-conjugated antibodies, according to the manufacturer’s instructions. All FACS/sorting and AbSeq antibodies used in this study are listed in **Table S1**.

### CD80/86 immunophenotyping

Immunophenotyping of the co-stimulatory molecules CD80/CD86 and CTLA-4 was performed in freshly isolated PBMCs. Cells were initially stained with fluorochrome-conjugated antibodies against surface receptors (see **Table S1**) in BD Brilliant stain buffer (BD Biosciences) for 30 min at room temperature. Fixation and permeabilisation was performed using FOXP3 Fix/Perm Buffer Set (eBioscience) according to the manufacturer’s instructions, and cells were then immunostained with fluorochrome-conjugated antibodies against intracellular markers (including CTLA-4, CD80 and CD86 where indicated) in BD Brilliant stain buffer for 45 min at room temperature. Immunostained samples were acquired using a BD Fortessa (BD Biosciences) flow cytometer with FACSDiva software (BD Biosciences) and analysed using FlowJo (Tree Star, Inc.).

### Tissue dissociation and isolation of CD45^+^ immune cells

For the characterisation of CD45^+^ immune cells from tissue and tissue, blood-derived PBMCs and paired duodenal biopsies were cryopreserved in CryoStor CS10 reagent (StemCell) and stored in liquid nitrogen until sample processing. Blood-derived PBMCs were processed as described above. The paired duodenal biopsies were thawed at 37°C in X-Vivo with 1% heat-inactivated, filtered human AB serum then subjected to gentle mechanical dissociation using gentleMACS (Miltenyi Biotec) followed by short 20 min enzymatic dissociation at 37°C using a very low concentration of Liberase TL (0.042 mg/ml; Sigma), 10 nM HEPES and 1mg/ml DNase I in X-Vivo with 10% FBS. Following enzymatic dissociation, the biopsies were homogenised using a more vigorous gentleMACS cycle and strained through a 70um filter with physical maceration to generate single-cell suspensions. CD45^+^ immune cells were further enriched using a 70/35% Percoll gradient (Sigma). The dissociation protocol and low concentration of Liberase TL enzyme were optimised to show minimal effect on the degradation of surface protein expression levels, as assessed by flow cytometry. This was critical to ensure maximal sensitivity and specificity of the AbSeq protein quantification in these samples.

Blood- and tissue-derived single-cell suspensions were incubated with Fixable Viability Dye eFluor 780 for 15 min at room temperature and total CD45^+^ cells were isolated by FACS sorting. Following FACS sorting the individual blood and tissue-derived subsets were incubated with Fc block reagent (BD Biosciences) and Sample Tag antibodies for 20 min at room temperature. Following three rounds of washing, cells were counted and equal numbers (35,000 cells) of blood- and tissue-derived cells from the same donor were pooled together and incubated with AbSeq antibody mastermix (**Table S1**) according to the manufacturer’s instructions. Cells were then washed two times in cold sample buffer, counted and resuspended in 620 ul of cold sample Buffer at a final concentration of 40 cells/ul for loading on a BD Rhapsody cartridge.

### cDNA library preparation and sequencing

Single-cell capture and cDNA library preparation was performed using the BD Rhapsody Express Single-cell analysis system (BD Biosciences), according to the manufacturer’s instructions. Briefly, cDNA was amplified - 10 cycles for resting cells and 9 cycles for *in vitro* activated cells - using the Human Immune Response primer panel (BD Biosciences), containing 399 primer pairs, targeting 397 different genes. The resulting PCR1 products were purified using AMPure XP magnetic beads (Beckman Coulter) and the respective mRNA and AbSeq/Sample tag products were separated based on size-selection, using different bead ratios (0.7X and 1.2X, respectively). The purified mRNA and Sample tag PCR1 products were further amplified (10 cycles), and the resulting PCR2 products purified by size selection (1X and 1.2X for the mRNA and Sample tag libraries, respectively). The concentration, size and integrity of the resulting PCR products was assessed using both Qubit (High Sensitivity dsDNA kit; Thermo Fisher) and the Agilent 4200 Tapestation system (High Sensitivity D1000 screentape; Agilent). The final products were normalised to 2.5 ng/ul (mRNA), 0.5 ng/ul (Sample tag) and 0.275 ng/ul (AbSeq) and underwent a final round of amplification (6 cycles for mRNA and 8 cycles for Sample Tag and AbSeq) using indexes for Illumina sequencing to prepare the final libraries. Final libraries were quantified using Qubit and Agilent Tapestation and pooled (∼38/58/2% mRNA/AbSeq/Sample tag ratio) to achieve a final concentration of 5nM. Final pooled libraries were spiked with 10% PhiX control DNA to increase sequence complexity and sequenced (75bp paired-end) on HiSeq 4000 sequencer (Illumina).

### Data analysis and QC

The FASTQ files obtained from sequencing were analysed following the BD Biosciences Rhapsody pipeline (BD Biosciences). Initially, read pairs with low quality are removed based on read length, mean base quality score and highest single nucleotide frequency. The remaining high-quality R1 reads are analysed to identify cell label and unique molecular identifier (UMI) sequences. The remaining high-quality R2 reads are aligned to the reference panel sequences (mRNA and AbSeq) using Bowtie2. Reads with the same cell label, same UMI sequence and same gene are collapsed into a single molecule. The obtained counts are adjusted by BD Biosciences developed error correction algorithms – recursive substitution error correction (RSEC) and distribution-based error correction (DBEC) – to correct sequencing and PCR errors. Cell counts are then estimated, using second derivative analysis to filter out noise cell labels, based on the assumption that putative cells have much more reads than noise cell labels. Thus when cells are sorted in the descending order by number of reads, the inflection point can be observed on a log transformed cumulative curve of number of reads. For the CD45^+^ sorted cells, due to the heterogeneity of the sample, we observed two inflection points (and two corresponding second derivative minima), and therefore, only cell labels after the second inflection point were considered noise labels. Barcoded oligo-conjugated antibodies (single-cell multiplexing kit; BD Biosciences) were used to infer origin of sample (ie. sorted cell population) and multiplet rate by the BD Rhapsody analysis pipeline.

The DBEC-adjusted molecule counts obtained from the Rhapsody pipeline were imported and the expression matrices were further analysed using R package Seurat 3.0 [42]. Most cells identified as undetermined by the Rhapsody pipeline had low number of features (mRNA and protein reads). These cells along with other cells with similarly low (<35) number of features were filtered out. Identified multiplet cells were also filtered out at this stage. A detailed summary of the number of putative captured cells, estimated multiplet rate and number of cells filtered from the analysis in each of the three experiments performed in this study is provided in **table S2**. The resulting matrices were log normalised using the default parameters in Seurat and the UMI counts were regressed out when scaling data. Uniform Manifold Approximation and Projection (UMAP) was used for dimensionality reduction. The default number of used dimensions of PCA reduction was increased to 30 based on Seurat elbow plot. For clustering, we increased the default resolution parameter value for clustering to 1.2 to obtain more fine-grained set of clusters. Differential expression analysis was performed using negative binomial generalized linear model implemented in Seurat, and integration of data from multiple experiments was performed using a combination of canonical correlation analysis (CCA) and identification of mutual nearest neighbours (MNN), implemented in Seurat 3.0 [43].

The Seurat objects were further converted and imported to the SCANPY toolkit [44] for consecutive analyses. We have computed diffusion pseudotime according to Haghverdi L et al. [45] which is implemented within SCANPY and used the Partition-based graph abstraction (PAGA) method [18] for formal trajectory inference and to detect differentiation pathways. For visualisation purposes we discarded low-connectivity edges using the threshold of 0.7. Additionally we have also performed pseudotime analysis using another independent method: Single-Cell Trajectories Reconstruction (STREAM) [20]. In this case to generate appropriate input files the Seurat objects were subsampled to include N=2,500 cells. The values other parameters not mentioned here were set to default.

## Abbreviations

CD: coeliac disease
FACS: Fluorescence-activated cell sorting
FC: fold-change
scRNA-seq: single-cell RNA-sequencing
SLE: systemic lupus erythematosus
T1D: type 1 diabetes
Teff: effector T cell
Treg: CD4^+^ regulatory T cell
UMAP: Uniform Manifold Approximation and Projection
UMI: unique molecular identifier

## Declarations

### Ethics approval and consent to participate

All samples and information were collected with written and signed informed consent. The study was approved by the local Peterborough and Fenland research ethics committee (05/Q0106/20). The D-GAP study was approved by the Royal Free Hospital & Medical School research ethics committee; REC (08/H0720/25). Small bowel biopsies were collected as part of a routine gastroduodenoscopy. Consent for research was obtained via the Oxford GI illnesses biobank (REC 16/YH/0247).

## Acknowledgments

We thank donors and patients for participation in this study. We gratefully acknowledge the participation of all NIHR Cambridge BioResource volunteers, and thank the NIHR Cambridge BioResource centre and staff for their contribution. We thank C. Guy from the Department of Paediatrics, University of Cambridge for D-GAP sample recruitment. We thank the volunteers from the Oxford Biobank (www.oxfordbiobank.org.uk) for their participation in this recall study. The Oxford BioBank and Oxford Bioresource are funded by the NIHR Oxford Biomedical Research Centre (BRC). We acknowledge the contribution of the Oxford Gastrointestinal biobank, which is supported by the NIHR Oxford Biomedical Research Centre. We also thank Georgina Burdon, Shannah Donhou, Sarune Kacinskaite and Heather McMurray, University of Oxford for sample collection and preparation and members of the Diabetes and Inflammation Laboratory for critical discussion.

## Supplementary Materials

**Table S1.** FACS and AbSeq anti-human monoclonal antibodies used in this study.

**Table S2.** Summary of the cell capture efficiency and multiplet rates for the experiments performed in this study

**Fig. S1.**
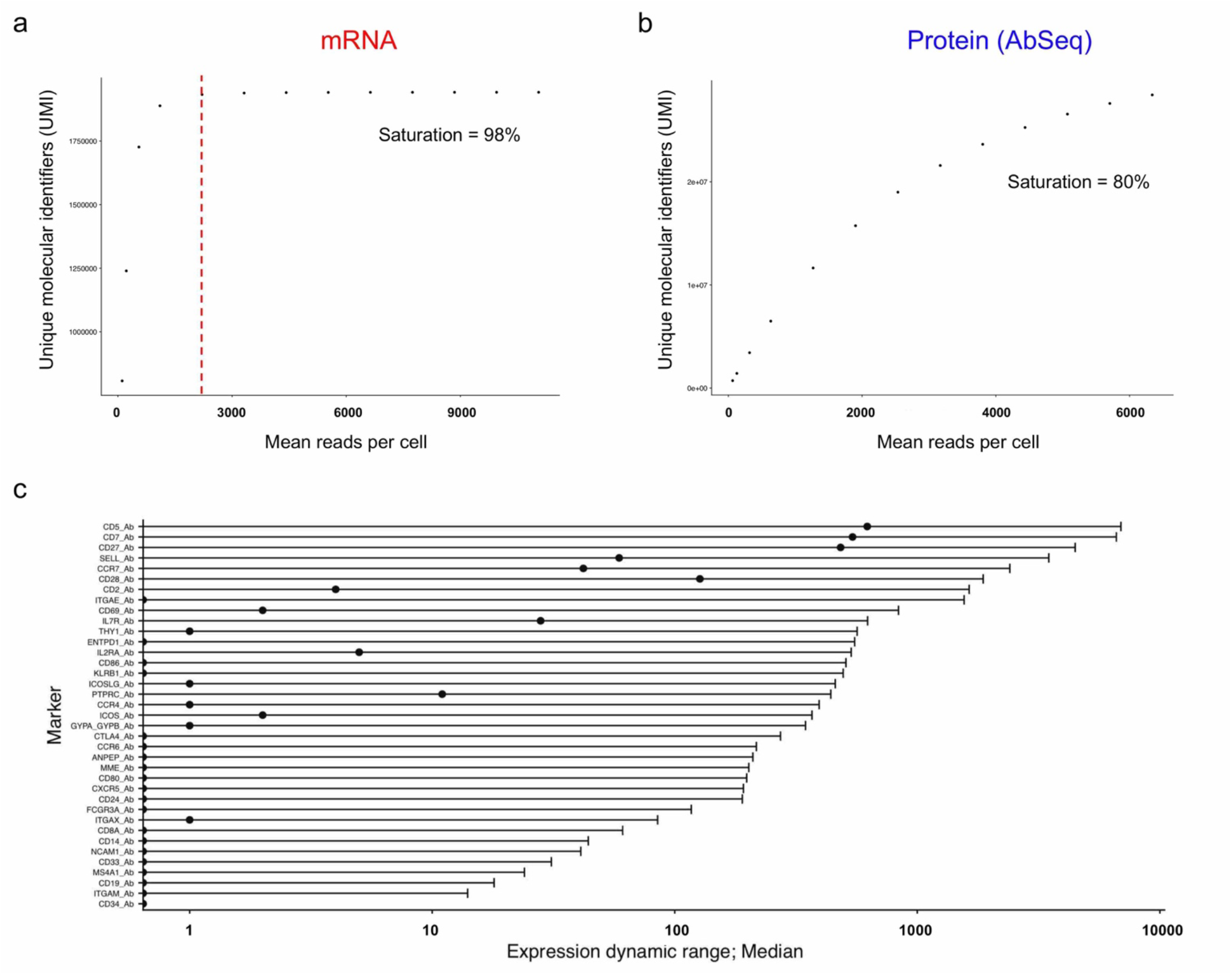
Protein expression displays much larger dynamic range of expression. (**a, b**) Sequencing saturation metrics of the mRNA (**a**) and protein (**b**) libraries. Saturation of the sequencing libraries was quantified as the number of identified distinct, non-clonally amplified cDNA molecules, marked by a unique molecular identifier (UMI), with increasing sequencing coverage. (**c**) Data shown depicts the distribution (median and range) of the expression of the 42 protein targets measured by AbSeq. Data was derived from the analysis of the first experiment performed on pre-sorted resting CD4^+^ T cell populations from a systemic lupus erythematosus (SLE) patient.

**Fig. S2.**
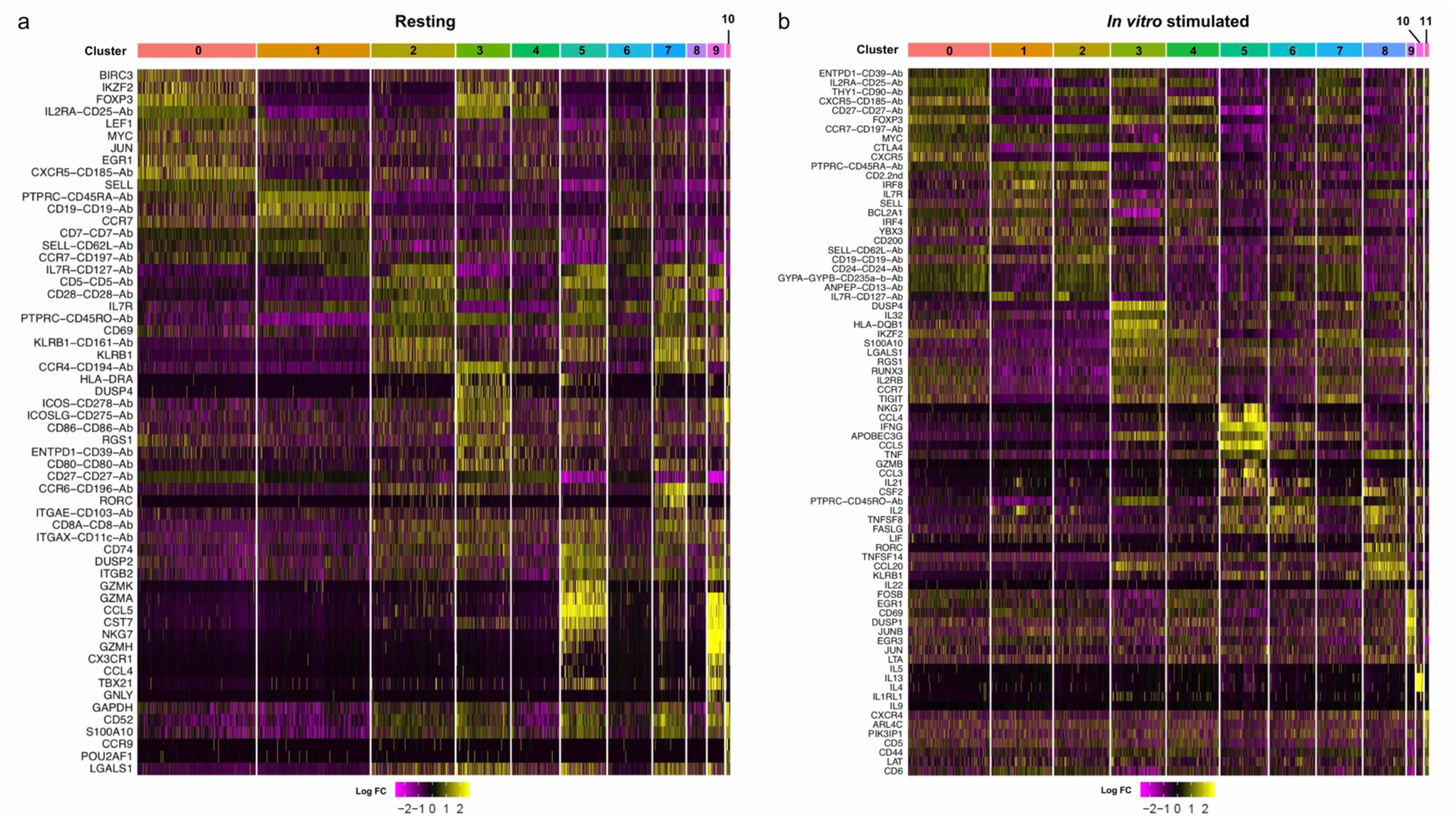
Differential expression in the identified resting and *in vitro* stimulated primary CD4^+^ T-cell subsets. (**a, b**) Heatmaps displaying the top 10 differentially expressed genes in each identified resting (**a**) or *in vitro* stimulated (**b**) CD4^+^ T-cell clusters. Stimulation condition involved a short period of incubation (90 min) with PMA + ionomycin.

**Fig. S3.**
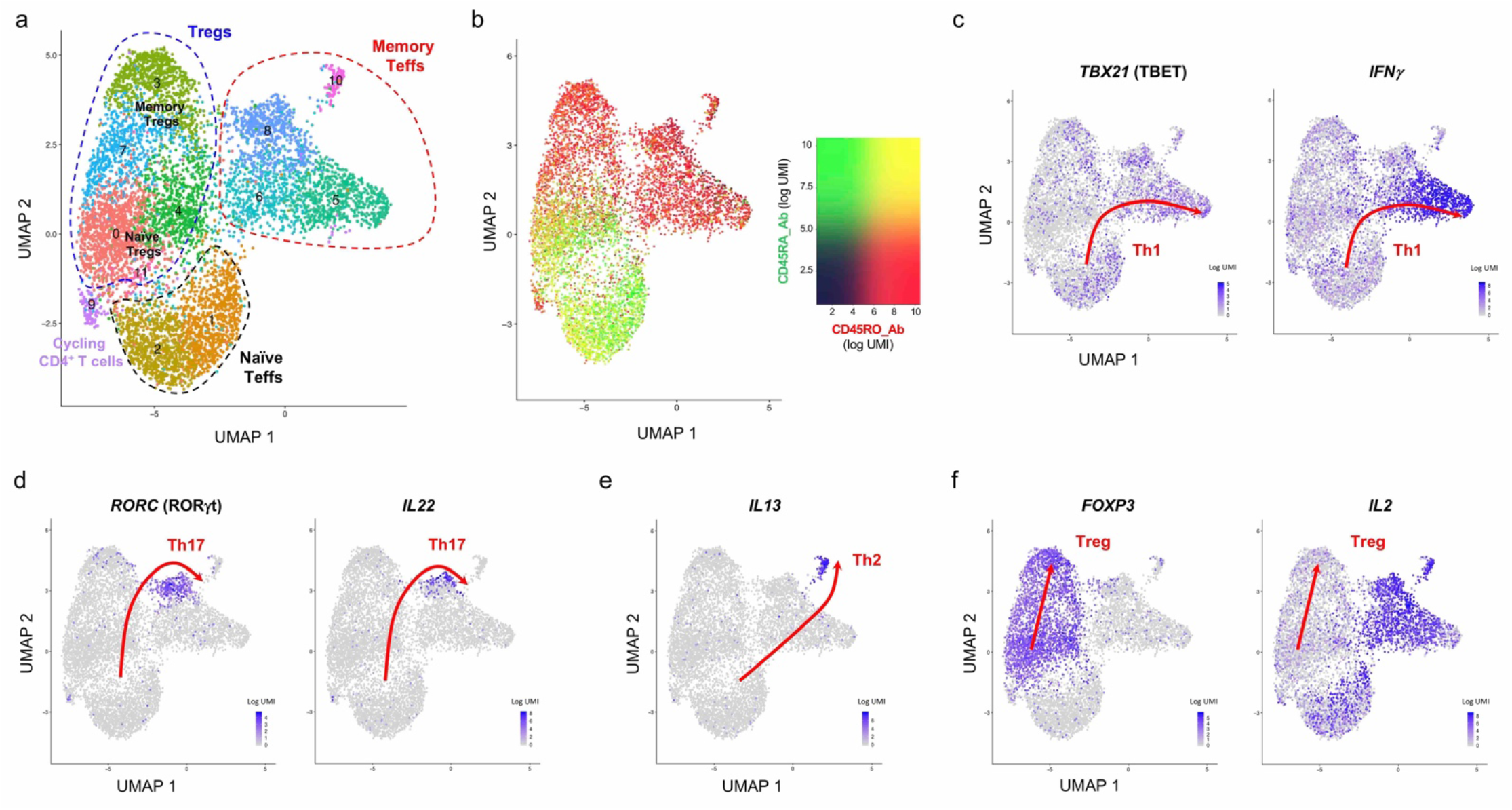
*In vitro* stimulation reinforces the trajectories of CD4^+^ T cell differentiation. **(a)** UMAP plot depicting the clustering of the *in vitro* stimulated primary CD4^+^ T cells (n = 7,265) isolated from blood of a systemic lupus erythematosus (SLE) patient. Stimulation condition involved a short period of incubation (90 min) with PMA + ionomycin. (**b**) Data shown depicts the overlaid protein expression levels of the CD45RA (black to green) and CD45RO (black to red) isoforms in each CD4^+^ T cell following *in vitro* activation with PMA + ionomycin. (**c, d**) UMAP plots depicting the co-expression of the CD4^+^ T-cell lineage-defining CD4^+^ Th1 transcription factor TBET and the Th1 effector cytokine IFN-g (**c**), as well as the Th17 transcription factor RORgt and the Th17 effector cytokine IL-22 (**d**) after *in vitro* stimulation with PMA + ionomycin for 90 min. (**e**) Expression of the canonical Th2 effector molecule IL-13 in *in vitro* stimulated CD4^+^ T cells. (**f**) UMAP plot depicting the expression of the Treg transcription factor FOXP3 and the prototypical CD4^+^ Th1 Teff cytokine IL-2 in the identified CD4^+^ T-cell subsets after *in vitro* stimulation with PMA + ionomycin for 90 min.

**Fig. S4.**
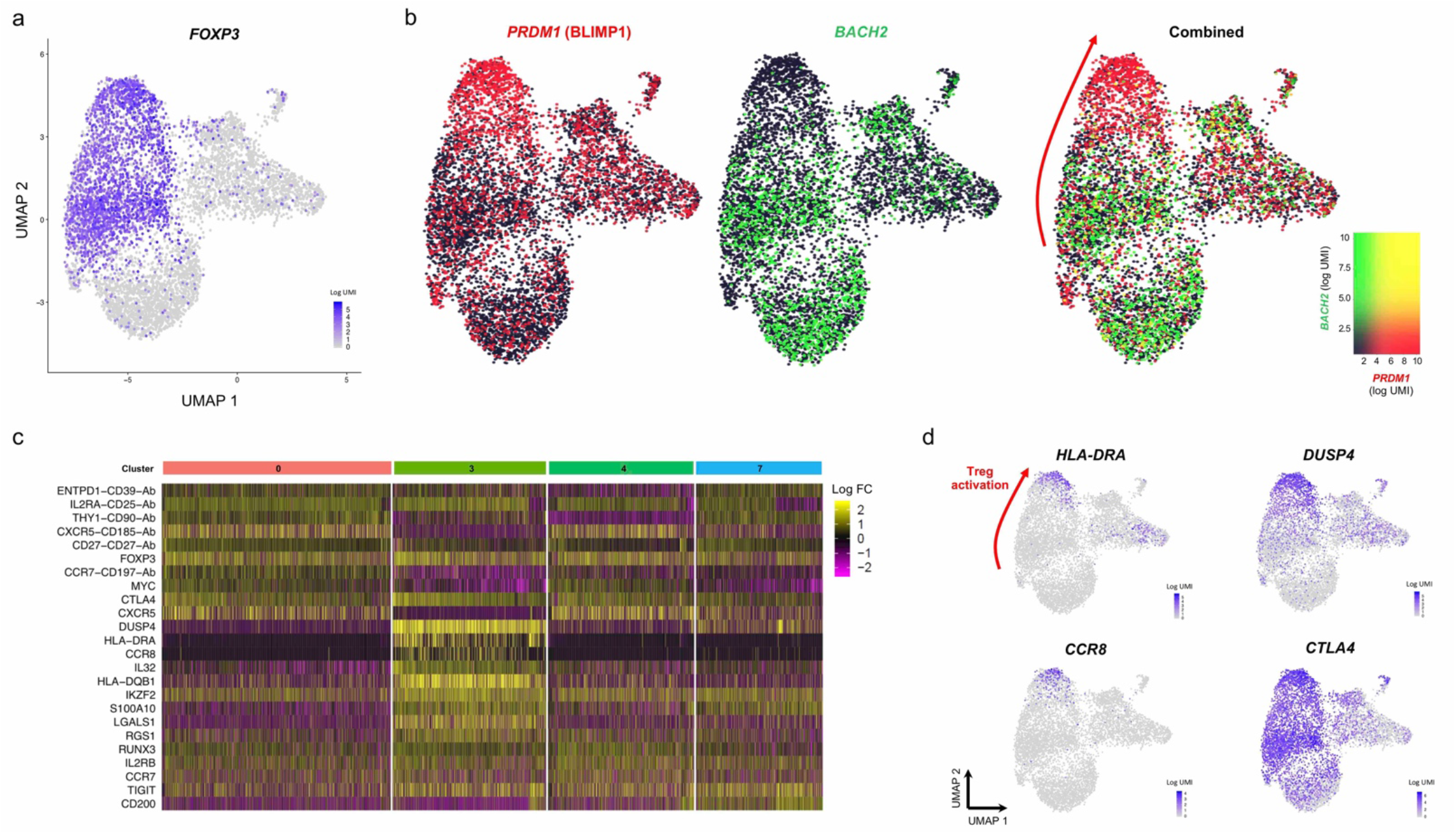
Interplay between the BACH2 and BLIMP-1 transcriptional programmes regulate CD4^+^ Treg activation in humans. (**a**) UMAP plot depicting the expression of the canonical Treg transcription factor FOXP3 in the identified *in vitro* stimulated CD4^+^ T-cell clusters. (**b**) Data shown depicts the overlaid protein expression levels of the transcription factors BACH2 (black to green) and BLIMP-1 (encoded by *PRDM1*; black to red) in each CD4^+^ T cell following *in vitro* activation with PMA + ionomycin. (**c**) Heatmap displaying the top 10 differentially expressed genes within the four identified Treg clusters following *in vitro* stimulation with PMA + ionomycin. (**d**) UMAP plots depicting the gradient of expression of highly differentially expressed genes in the cluster of activated Tregs (cluster 3) following *in vitro* stimulation with PMA + ionomycin. Red arrows in this figure indicate the gradient of decreasing *BACH2* and concomitant gain in *PRDM1* expression associated with the gradual expression of Treg activation molecules and the acquisition of an activated Treg phenotype.

**Fig. S5.**
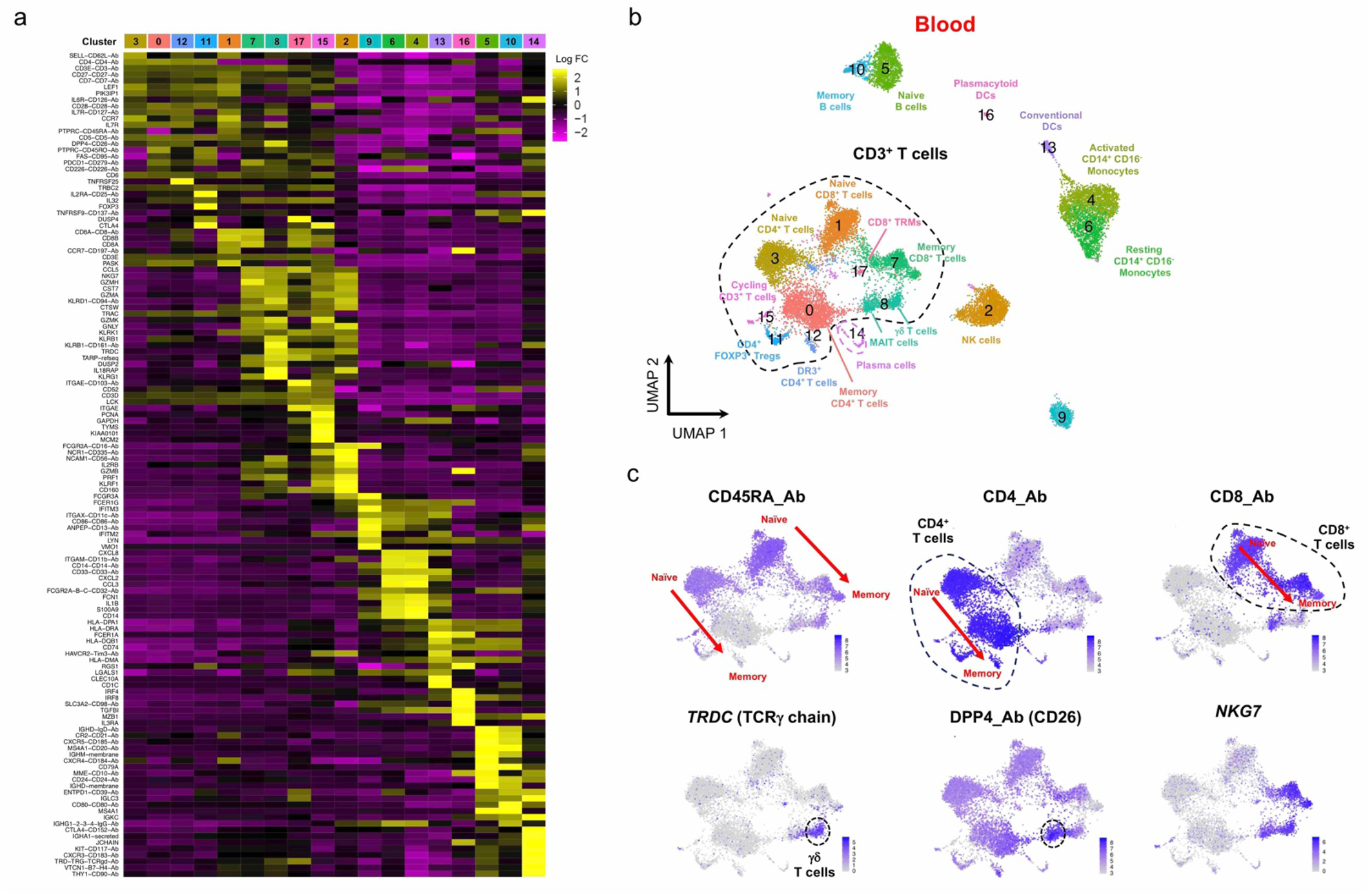
Single-cell mRNA and protein quantification identifies distinct functional populations of human circulating CD3^+^ T cells. (**a**) Heatmap displaying the top 10 differentially expressed genes in each identified cluster from the CD45^+^ immune cells isolated from blood of two coeliac disease (CD) patients with active disease. (**b**) UMAP plot depicting the clustering of the circulating CD45^+^ immune cells. Dashed lines outline the annotated CD3^+^ T-cell clusters annotated from the differentially expressed genes in those clusters. (**c**) Functional annotation of the peripheral T-cell subsets using the expression profile of CD45RA and additional key lineage-defining T-cell markers such as CD4, CD8, *TRDC*, DPP4 and the effector cytokine gene *NKG7*. Arrows indicate the gradient of decreasing CD45RA and concomitant gain in CD45RO expression associated with the acquisition of a memory phenotype in response to antigen stimulation in CD4^+^ and CD8^+^ T-cells. DR3, death-receptor 3 (encoded by *TNFRSF25*); TRM, tissue-resident memory T cells; MAIT, mucosal-associated invariant T cells; ILC3, type 3 innate lymphoid cell.

**Fig. S6.**
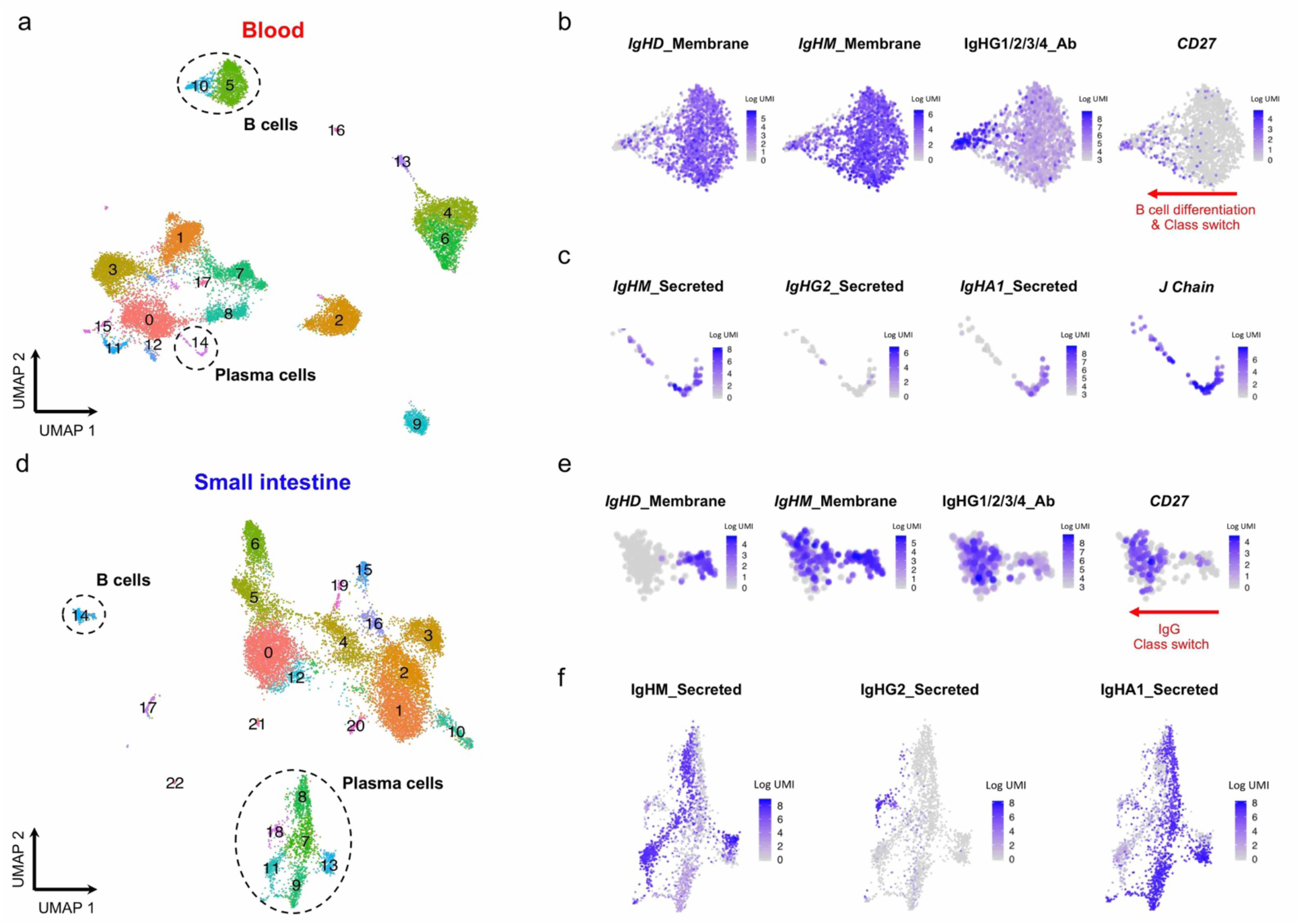
Targeted multi-omics approach reveals trajectories of B-cell differentiation and class switching in blood and tissue. (**a**) UMAP plots depicting the clustering of the total CD45^+^ immune cells isolated from blood of two coeliac disease (CD) patients with active disease. Dashed lines outline the annotated B-cell and plasma cell subsets annotated from the differentially expressed genes in those clusters. (**b, c**) Expression of key B-cell differentiation and class switching (**b**) and plasma cell (**c**) markers, including *CD27* mRNA and selected surface expressed or secreted immunoglobulin (Ig) receptors in the identified circulating B-cell and plasma cell clusters. (**d**) UMAP plots depict the clustering of the total CD45^+^ immune cells isolated from duodenal tissue biopsies. Dashed lines outline the annotated B-cell and plasma cell subsets annotated from the differentially expressed genes in those clusters. (**e, f**) Expression of key B-cell differentiation and class switching (**e**) and plasma cell (**f**) markers, including *CD27* mRNA and selected surface expressed or secreted immunoglobulin (Ig) receptors in the identified in the tissue-resident B-cell and plasma cell clusters.

## References

1. Ornatsky O, Bandura D, Baranov V, Nitz M, Winnik MA, Tanner S. Highly multiparametric analysis by mass cytometry. J Immunol Methods. 2010;361:1–20.

2. See P, Lum J, Chen J, Ginhoux F. A Single-Cell Sequencing Guide for Immunologists. Front. Immunol. 2018; 2425.

3. Papalexi E, Satija R. Single-cell RNA sequencing to explore immune cell heterogeneity. Nat Rev Immunol. 2017;18:35.

4. Picelli S, Björklund ÅK, Faridani OR, Sagasser S, Winberg G, Sandberg R. Smart-seq2 for sensitive full-length transcriptome profiling in single cells. Nat Methods. 2013;10:1096.

5. Zheng GXY, Terry JM, Belgrader P, Ryvkin P, Bent ZW, Wilson R, et al. Massively parallel digital transcriptional profiling of single cells. Nat Commun. 2017;8:14049.

6. Gierahn TM, Wadsworth II MH, Hughes TK, Bryson BD, Butler A, Satija R, et al. Seq-Well: portable, low-cost RNA sequencing of single cells at high throughput. Nat Methods. 2017;14:395.

7. Macosko EZ, Basu A, Satija R, Nemesh J, Shekhar K, Goldman M, et al. Highly Parallel Genome-wide Expression Profiling of Individual Cells Using Nanoliter Droplets. Cell. 2015;161:1202–14.

8. Andrews TS, Hemberg M. False signals induced by single-cell imputation. F1000Research. 2018;7.

9. Stoeckius M, Hafemeister C, Stephenson W, Houck-Loomis B, Chattopadhyay PK, Swerdlow H, et al. Simultaneous epitope and transcriptome measurement in single cells. Nat Methods. 2017;14:865.

10. Peterson VM, Zhang KX, Kumar N, Wong J, Li L, Wilson DC, et al. Multiplexed quantification of proteins and transcripts in single cells. Nat Biotechnol. 2017;35:936.

11. Shahi P, Kim SC, Haliburton JR, Gartner ZJ, Abate AR. Abseq: Ultrahigh-throughput single cell protein profiling with droplet microfluidic barcoding. Sci Rep. 2017;7:44447.

12. Ferreira RC, Simons HZ, Thompson WS, Rainbow DB, Yang X, Cutler AJ, et al. Cells with Treg-specific FOXP3 demethylation but low CD25 are prevalent in autoimmunity. J Autoimmun. 2017;84:75–86.

13. Stoeckius M, Zheng S, Houck-Loomis B, Hao S, Yeung BZ, Mauck WM, et al. Cell Hashing with barcoded antibodies enables multiplexing and doublet detection for single cell genomics. Genome Biol. 2018;19:224.

14. McInnes L, Healy J, Melville J. UMAP: Uniform Manifold Approximation and Projection for Dimension Reduction. arXiv. 2018;arXiv:1802.

15. Sallusto F, Lenig D, Förster R, Lipp M, Lanzavecchia A. Two subsets of memory T lymphocytes with distinct homing potentials and effector functions. Nature. 1999;401:708–12.

16. Zemmour D, Zilionis R, Kiner E, Klein AM, Mathis D, Benoist C. Single-cell gene expression reveals a landscape of regulatory T cell phenotypes shaped by the TCR. Nat Immunol. 2018;19:291–301.

17. Miragaia RJ, Gomes T, Chomka A, Jardine L, Riedel A, Hegazy AN, et al. Single-Cell Transcriptomics of Regulatory T Cells Reveals Trajectories of Tissue Adaptation. Immunity. 2019;50:493–504.e7.

18. Wolf FA, Hamey FK, Plass M, Solana J, Dahlin JS, Göttgens B, et al. PAGA: graph abstraction reconciles clustering with trajectory inference through a topology preserving map of single cells. Genome Biol. 2019;20:59.

19. Weaver CT, Hatton RD. Interplay between the TH17 and TReg cell lineages: a (co-) evolutionary perspective. Nat Rev Immunol. 2009;9:883–9.

20. Chen H, Albergante L, Hsu JY, Lareau CA, Lo Bosco G, Guan J, et al. Single-cell trajectories reconstruction, exploration and mapping of omics data with STREAM. Nat Commun. 2019;10:1903.

21. Walker LSK, Sansom DM. The emerging role of CTLA4 as a cell-extrinsic regulator of T cell responses. Nat Rev Immunol. 2011;11:852.

22. Dopfer EP, Minguet S, Schamel WWA. A New Vampire Saga: The Molecular Mechanism of T Cell Trogocytosis. Immunity. 2011;35:151–3.

23. Schubart DB, Rolink A, Kosco-Vilbois MH, Botteri F, Matthias P. B-cell-specif ic coactivator OBF-1/OCA-B/Bob1 required for immune response and germinal centre formation. Nature. 1996;383:538–42.

24. Shakya A, Goren A, Shalek A, German CN, Snook J, Kuchroo VK, et al. Oct1 and OCA-B are selectively required for CD4 memory T cell function. J Exp Med. 2015;212:2115– 2131.

25. Stauss D, Brunner C, Berberich-Siebelt F, Höpken UE, Lipp M, Müller G. The transcriptional coactivator Bob1 promotes the development of follicular T helper cells via Bcl6. EMBO J. 2016;35:881–98.

26. Kumar B V, Ma W, Miron M, Granot T, Guyer RS, Carpenter DJ, et al. Human Tissue-Resident Memory T Cells Are Defined by Core Transcriptional and Functional Signatures in Lymphoid and Mucosal Sites. Cell Rep. 2017;20:2921–34.

27. Szabo PA, Levitin HM, Miron M, Snyder ME, Senda T, Yuan J, et al. A single-cell reference map for human blood and tissue T cell activation reveals functional states in health and disease. bioRxiv. 2019;555557.

28. Stubbington MJT, Rozenblatt-Rosen O, Regev A, Teichmann SA. Single-cell transcriptomics to explore the immune system in health and disease. Science. 2017;358:58–63.

29. Tanay A, Regev A. Scaling single-cell genomics from phenomenology to mechanism. Nature. 2017;541:331.

30. Mimitou EP, Cheng A, Montalbano A, Hao S, Stoeckius M, Legut M, et al. Multiplexed detection of proteins, transcriptomes, clonotypes and CRISPR perturbations in single cells. Nat Methods. 2019;16:409–12.

31. Gerlach JP, van Buggenum JAG, Tanis SEJ, Hogeweg M, Heuts BMH, Muraro MJ, et al. Combined quantification of intracellular (phospho-)proteins and transcriptomics from fixed single cells. Sci Rep. 2019;9:1469.

32. Gu P, Fang Gao J, D’Souza CA, Kowalczyk A, Chou K-Y, Zhang L. Trogocytosis of CD80 and CD86 by induced regulatory T cells. Cell Mol Immunol. 2012;9:136.

33. Brown R, Kabani K, Favaloro J, Yang S, Ho PJ, Gibson J, et al. CD86+ or HLA-G+ can be transferred via trogocytosis from myeloma cells to T cells and are associated with poor prognosis. Blood. 2012;120:2055–2063.

34. Ovcinnikovs V, Ross EM, Petersone L, Edner NM, Heuts F, Ntavli E, et al. CTLA-4– mediated transendocytosis of costimulatory molecules primarily targets migratory dendritic cells. Sci Immunol. 2019;4:eaaw0902.

35. Huang Y, Li Y, Wei B, Wu W, Gao X. CD80 Regulates Th17 Cell Differentiation in Coxsackie Virus B3-Induced Acute Myocarditis. Inflammation. 2018;41:232–9.

36. Constant S, Pfeiffer C, Woodard A, Pasqualini T, Bottomly K. Extent of T cell receptor ligation can determine the functional differentiation of naive CD4+ T cells. J Exp Med. 1995;182:1591–1596.

37. Brogdon JL, Leitenberg D, Bottomly K. The Potency of TCR Signaling Differentially Regulates NFATc/p Activity and Early IL-4 Transcription in Naive CD4+ T Cells. J Immunol. 2002;168:3825–3832.

38. Höllsberg P, Scholz C, Anderson DE, Greenfield EA, Kuchroo VK, Freeman GJ, et al. Expression of a hypoglycosylated form of CD86 (B7-2) on human T cells with altered binding properties to CD28 and CTLA-4. J Immunol. 1997;159:4799–4805.

39. Caielli S, Veiga DT, Balasubramanian P, Athale S, Domic B, Murat E, et al. A CD4+ T cell population expanded in lupus blood provides B cell help through interleukin-10 and succinate. Nat Med. 2019;25:75–81.

40. Schwanhäusser B, Busse D, Li N, Dittmar G, Schuchhardt J, Wolf J, et al. Global quantification of mammalian gene expression control. Nature. 2011;473:337.

41. Vogel C, Marcotte EM. Insights into the regulation of protein abundance from proteomic and transcriptomic analyses. Nat Rev Genet. 2012;13:227.

42. Butler A, Hoffman P, Smibert P, Papalexi E, Satija R. Integrating single-cell transcriptomic data across different conditions, technologies, and species. Nat Biotechnol. 2018;36:411.

43. Stuart T, Butler A, Hoffman P, Hafemeister C, Papalexi E, Mauck WM, et al. Comprehensive Integration of Single-Cell Data. Cell. 2019;177:1888–1902.e21.

44. Wolf FA, Angerer P, Theis FJ. SCANPY: large-scale single-cell gene expression data analysis. Genome Biol. 2018;19:15.

45. Haghverdi L, Büttner M, Wolf FA, Buettner F, Theis FJ. Diffusion pseudotime robustly reconstructs lineage branching. Nat Methods. 2016;13:845.

